# GliaMorph: A modular image analysis toolkit to quantify Müller glial cell morphology

**DOI:** 10.1101/2022.05.05.490765

**Authors:** Elisabeth Kugler, Isabel Bravo, Xhuljana Durmishi, Stefania Marcotti, Sara Beqiri, Alicia Carrington, Brian M. Stramer, Pierre Mattar, Ryan B. MacDonald

**Affiliations:** Institute of Ophthalmology, University College London, 11-43 Bath St, Greater London EC1V 9EL.; Randall Centre for Cell & Molecular Biophysics, King’s College London, New Hunt’s House, London SE1 1UL.; Department of Cellular and Molecular Medicine, University of Ottawa, Ottawa, ON K1H 8M5, Canada.; Ottawa Hospital Research Institute (OHRI), Ottawa, ON K1H 8L6, Canada.

**Keywords:** retina, Müller glia, glia morphology, zebrafish, development

## Abstract

Cell morphology is critical for all cell functions. This is particularly true for glial cells as they rely on their complex shape to contact and support neurons. However, methods to quantify complex glial cell shape accurately and reproducibly are lacking. To address this gap in quantification approaches, we developed an analysis pipeline called “*GliaMorph*”. GliaMorph is a modular image analysis toolkit developed to perform (i) image pre-processing, (ii) semi-automatic region-of-interest (ROI) selection, (iii) apicobasal texture analysis, (iv) glia segmentation, and (v) cell feature quantification. Müller Glia (MG) are the principal retinal glial cell type with a stereotypic shape linked to their maturation and physiological status. We here characterized MG on three levels, including (a) global image-level, (b) apicobasal texture, and (c) apicobasal vertical-to-horizontal alignment. Using GliaMorph, we show structural changes occurring in the developing retina. Additionally, we study the loss of *cadherin2* in the zebrafish retina, as well as a glaucoma mouse disease model. The GliaMorph toolkit enables an in-depth understanding of MG morphology in the developing and diseased retina.

**Graphical Abstract:** 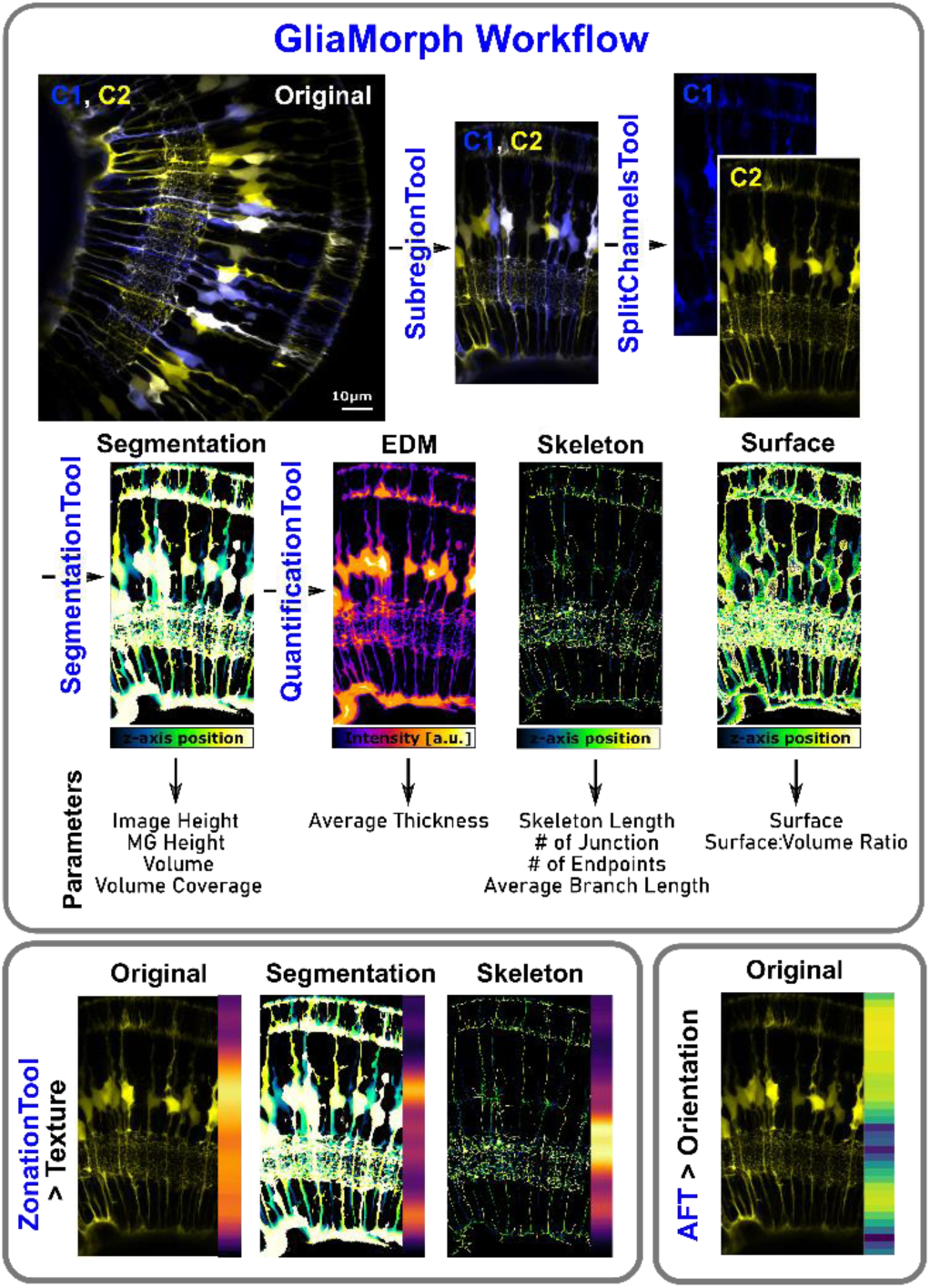

**Highlights:**

- Glial morphology is complex, making it challenging to accurately quantify 3D cell shape.
- We developed the GliaMorph toolkit for image pre-processing, glial segmentation, and quantification of Müller glial cells.
- Müller glia elaborate their morphology and rearrange subcellular features during embryonic development.
- GliaMorph accurately identifies subcellular changes in models with disrupted glia cells, including zebrafish *cadherin2* loss of function and a mouse glaucoma model.

## Introduction

While a vast amount of biomedical research relies on microscopy data and image-driven research, methods to objectively as well as reproducibly process cell morphology are lacking. However, computational analysis is paramount to understanding cell function and connectivity on a more abstract level, particularly in complex tissues. Glial cells are some of the most morphologically elaborate cells and provide a myriad of functions in the central nervous system (CNS) [1], [2]. To fulfil these critical functions glial cells are precisely shaped to contact neurons, their synapses, and/or the vasculature. Glial shape is not only pivotal in healthy tissue, but is altered in numerous neurodegenerative conditions, and can precede neuronal dysfunction in some cases such as epilepsy [3] or diabetic retinopathy [4]. Hence, measuring MG morphology in a robust and reliable manner is paramount to our understanding of CNS development and dysfunction. Current methods for glial cell morphological analysis often require user input (*i.e.*, manual cell tracing) that might result in subjective bias, offer crude measurements (*i.e.,* not sub-cellular resolution and not reproducible detail), or are challenging to adapt to specific biological questions and dynamic timelapse acquisitions. This leads to a data analysis bottleneck in morphological analysis and image-based cell profiling [5], [6]. As such, it is necessary to develop workflows and high-quality datasets of glial morphologies for robust and (semi-)automatic analysis of glial shape in healthy and diseased CNS.

Computationally, glia cell analysis is challenging as glia have highly complex morphologies [7], [8]. To analyze glia shape for (semi-)automatically, cells are extracted from images by an image binarization step, called segmentation. However, due to their elaborate shape, glial cells are more challenging to segment automatically than for example cells with a simple round shape, such as red blood cells. In addition to complex MG shape, most glia visualization techniques do not visualize glia cells consistently but suffer from heterogeneity, such as irregular antibody or fluorophore accumulation [9], [10]. This inconsistency makes it challenging to segment different glial subregions equally. Lastly, cells can be described by various different features, but the relevance and applicability of features depends on the biological question [6]. In the era of machine learning, defining relevant features could be assumed to be easily achievable. However, machine learning methods require appropriate training sets or *a priori* information. Therefore, conventional image analysis workflows are required to establish benchmark datasets before machine learning approaches could be considered. Hence, there is a rationale to develop computational (semi-)automatic analysis methods to resolve glial cell morphology and status [9].

As a part of the CNS, the retina serves as a tractable model to study due to highly stereotypic architecture. The retina consists of 7 main cell types, six neuronal and one glial cell type called Müller glia (MG). All cells are organized into stereotypic neuronal layers, creating retinal lamination, with well understood neuronal circuits [11]. MG are considered molecular and functional homologs to astrocytes [12], as they carry out numerous physiological roles to support neurons [13], [14] and will emanate elaborate fine projections to contact synapses [8]. Embryonically, nascent MG cells derive from retinal progenitor cells, beginning as simple radial cells. These then mature to morphologically elaborate cells with a highly branched morphology [7], [15]. Mature MG morphology has been divided into five specific subregions (see **Fig. 1C**): domains (1) and (2) consist of the apical regions making up the outer limiting membrane (OLM) and elaborating within the outer plexiform layer (OPL); domain (3) MG includes the cell bodies in the inner nuclear layer (INL); domain (4) encompasses the cellular protrusions weaving through the inner plexiform layer (IPL); and domain (5) consist of the basal end feet which contact the underlying vasculature and the inner limiting membrane [8]. In addition to this apicobasal specialization, MG are organized laterally to interact with each other and other cell types within the retina. Thereby intercalating cells in a so-called “tiled” fashion and contacting almost all cells [7], [8]. Thus, MG morphology is linked to their spatial and functional organization. Moreover, it was shown that MG shape is indicative of their maturity [7] and health [10]. For instance, neuronal tissue damage can elicit MG to undergo gliosis, a reactive state whereby their morphology and gene expression levels are drastically altered; this is observed in several diseases such as glaucoma [16]. Hence, thorough analysis and understanding of MG morphology are crucial to the complete understanding of the overall health and performance of retinal neurons.

**Figure 1.**
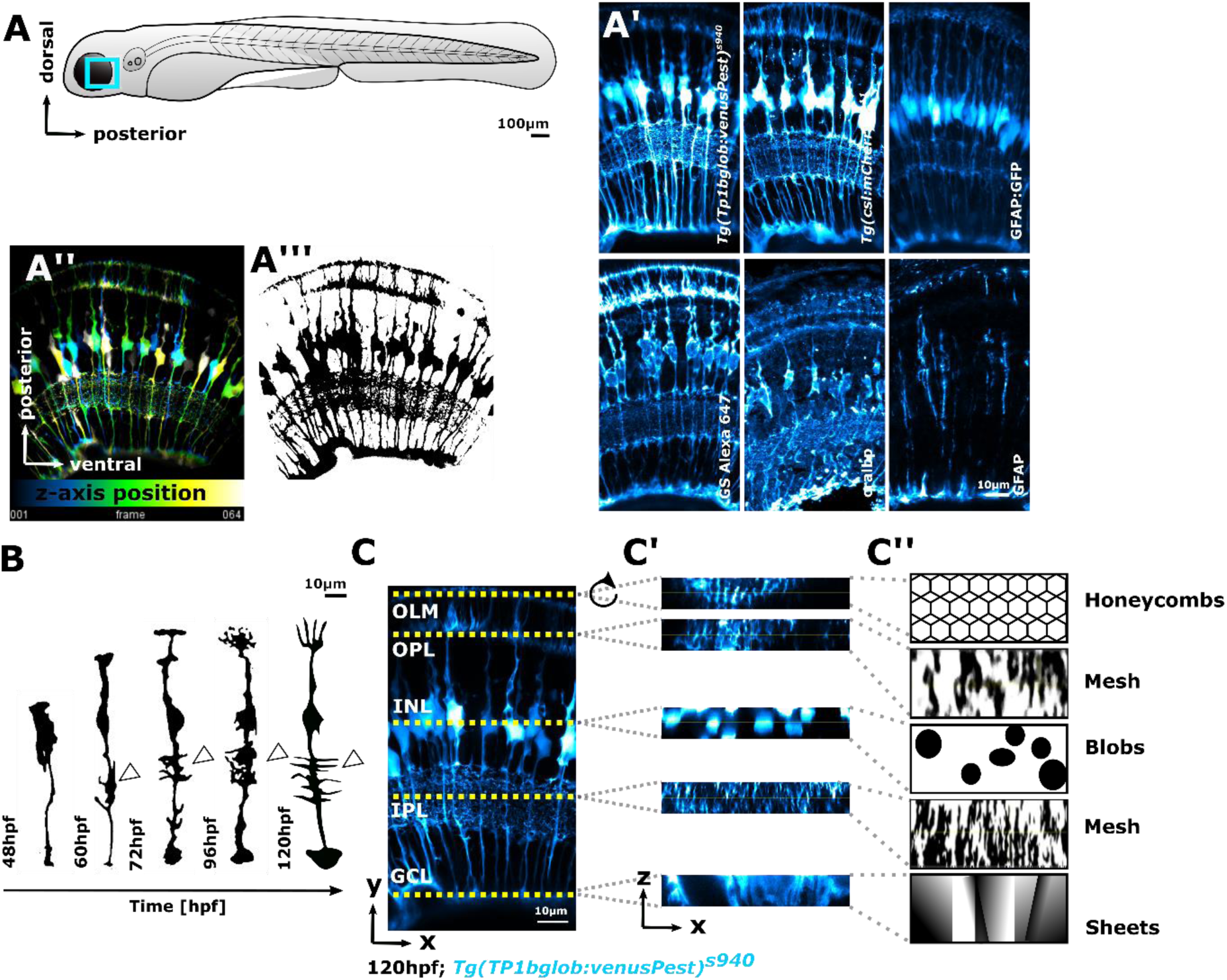
MG cells have a highly complex shape that can be visualized by specific cellular labels by confocal microscopy. **(A)** Imaging of MG was conducted in the ventro-temporal zebrafish retina to standardize the ROI. **(A’)** MG are the principal glia cell in the retina and can be visualized with transgenic reporter lines or immunohistochemistry stainings. **(A’’, A’’’)** Computationally, identifying individual MG cells is challenging as typically all MG cells are labelled by transgenic or antibody markers, making it a complex task to identify where individual cells start/end. **(B)** Schematic of individual MG cell morphological maturation during early development, showing elaboration of subregions and an increase in protrusions (arrowheads). **(C)** Maximum intensity projection (MIP) micrograph with yellow dotted lines indicating the apicobasal position of retinal layers. **(C’’)** Resliced/transformed section through the yellow dotted lines, visualizing that cell subregions have distinct properties along the apicobasal axis, making it challenging for computational approaches to be applied throughout the image in the same fashion. **(C’’)** Computational description of MG subregions features.

The overall organisation and composition of the retina, including MG function and structure, are highly conserved between zebrafish and humans [17], [18]. Further, zebrafish is a well-established model to study retina development and disease [17], [19]–[21]. The zebrafish retina is suitable for morphological analysis as it is accessible for advanced imaging, rapidly develops, and is amenable to various manipulation and/or visualization techniques. These include specific transgenic reporter lines [22] and antibodies labelling MG marker genes (e.g., Glutamine synthetase, GS) (**Fig. 1A)**. However, as these labels mark a considerable proportion of MGs in the retina, it is challenging, both morphologically and computationally, to identify or quantify individual cells (**Fig. 1A’’**). Furthermore, MG become increasingly morphologically elaborate during retinal development (**Fig. 1B**). Therefore, the zebrafish retina is suitable to generate the high-resolution imaging data required to develop a computational workflow to robustly quantify complex glial morphologies.

Here we establish a workflow, called “GliaMorph”, that allows for the reproducible assessment of glia shape. GliaMorph is a 3D image analysis toolkit to quantitatively describe the stereotypic cellular morphology of MG. Specifically, we present **(a)** an in-depth description of encountered data challenges, **(b)** steps for image pre-processing to improve data quality**; (c)** provide a novel tool for semi-automatic region-of-interest (ROI) selection to allow comparability between samples and groups, **(d)** a method to automatically plot apicobasal textures, **(e)** a MG segmentation workflow, and **(f)** a workflow for 3D quantification of glia. This allows for three-level image assessment, namely (a) global image-level, (b) apicobasal texture, and (c) apicobasal vertical-to-horizontal alignment. We apply this to retinas of (i) fully transgenic zebrafish, where all MG cells are labelled, and to (ii) single-cell transient injected clonal data, where individual MG cells are labelled. We show that MG become significantly more morphologically complex throughout their maturation 60 hours post fertilisation (hpf) 96 hpf. We also use CRISPR/Cas9 to manipulate MG morphogenesis at this critical stage. *Cadherin2* (*cdh2*) [23] crispants present MG morphological disruption in injected embryos, showing that GliaMorph is capable of detecting sub-cellular differences in MG shape in the developing zebrafish retina. Finally, we apply GliaMorph to immunohistochemistry data from the mouse glaucoma model DBA/2J [24] and detect morphological signs of gliosis, revealing this tool has the potential to work across species.

Taken together, our work presents a benchmark for 3D MG analysis across MG visualization techniques, developmental ages, and species. To allow the method dissemination, all steps were implemented in the open-source image analysis software Fiji [25] and the code, as well as example data, are provided.

## Results

### Data understanding for reverse experimental design and optimization

When developing image quantification approaches for cell morphology, data understanding is key for workflow development from a computational point of view. This is particularly important for understanding MG cell shape, as they have a complex apicobasal morphology (**Fig. 1A**). Here we use confocal imaging as it is a widely used imaging technique, allowing for sufficient resolution to resolve individual glial sub- domains [26]. The following sections will focus on how we optimized data acquisition and data quality. To standardize the ROI for image acquisition, we exclusively focus on the ventro-temporal retina, as regional differences in cell morphologies exist across the retina in zebrafish (**Fig 1A**) [27], [28]. MG have five precise subregions where each fulfils a specific function for nearby retinal neurons [7] (**Fig. 1C**). Computationally these regions fall into categories based on their geometries; *subregion 1* resembles a honeycomb structure, allowing MG to interact photoreceptors. *Subregion 2* resembles a fine mesh-like structure, allowing MG to interweave with synaptic terminal of photoreceptors, bipolar and horizontal cells. *Subregion 3* are blob-like cell bodies. *Subregion 4* are mesh-like protrusions in the IPL, and *subregion 5* are the sheet-like endfeet. As these subregions/domains become elaborate over time and are affected by disease, glia shape is a direct readout of cell status.

In addition to apicobasal shape differences, MG subregions also are computationally distinctive concerning levels and patterns of signal (**Fig. S1A**). This suggests that (a) either local image processing, specifically tailored to these zones, could outperform global image processing approaches due to apicobasal inhomogeneity; or (b) that image pre-processing steps are required to equalize signal across the 3D volume. This is particularly important, as it is known that confocal-type imaging can suffer from z-axis signal decay, due to light interaction with matter [29]. We thus assessed lateral intensity profiles (x,y) that showed a homogenous signal distribution (**Fig. S1B**). However, as expected, this was not the case for axial (z) intensity profiles (**Fig. S1C**), suggesting that the images are not homogeneous in 3D for signal. Next, we wanted to assess the impact of z-step sampling frequency on data quality as this is an often overlooked factors for sufficient data quality. Nyquist rate for the widely used transgenic reporter line *Tg(TP1bglob:VenusPest)^s940^* [30], which specifically labels MG in the retina [31], was assessed (see Material and Methods for details). Optimized image acquisition led to improved data quality and MG were more connected through the z-axis (**Fig. S1E**). Lastly, image artefacts due to sample motion, such as cardiac contraction and muscle movement, can impact 3D quantification results. Indeed, we found stripe artefacts (**Fig. S1F),** blurring (**Fig. S1G**), and inter-plane (**Fig. S1H**) motion artefacts to be present when performing *in vivo* imaging. However, those were minimal and intra-stack motion correction [32] could be applied for more severe cases, such as focus drift. Together, this data understanding allowed us to draft required image processing steps and optimize data acquisition.

### Establishing image comparability between samples within data groups

For robust computation of glial morphology, each image dataset must be processed identically to allow images to be compared within, and between groups, as well as between different laboratories, users, or microscopes. Several factors can introduce variability that may skew results. For example, whole-mount zebrafish imaging requires samples embedding, which could affect the acquisitions, resulting in imaging different areas of the retina. This is accompanied by the fact that there are no hard- set standards on sample orientation. Additionally, the above-mentioned z-axis signal decay and different z-stack sizes between samples make image comparisons challenging. Together, this means that typically only regionally meaningful data are acquired. In addition, fields of view (FOV) are often larger than needed and images cannot be directly compared to each other but require pre-processing to allow comparable ROIs between samples. We thus developed a semi-automatic approach, called the ***subregionTool***, which enables ROIs extraction based on manual line selection (**Fig. 2A**), which applies to right and left eyes *i.e.,* the reverse (**Fig. S2A,B**). Once the ROI is selected, the images are then aligned along the y-axis (**Fig. S2C,D**). Then a bounding box is created with the line at the centre (unless the image is too small; see **Fig. S2E,F,** and code for details) and the stacks are reduced to the specified depth (**Fig. S2G**). For cases where the image is rectangular and not cubic, rotation can result in decreased image height (**Fig. S2H**). To avoid this, rotation is suggested before application of the *subregionTool* (using the ***90DegreeRotation*** Macro). This *subregionTool* is also applicable to other imaging datasets, not only MG, such as retinal neurons. Hence, we were able to overlap data from different neuronal markers, as well as images from 24-to-72 hpf including neurons as well as MG (**Fig. 2B**). To confirm data comparability, we measured progenitor/MG height, showing high similarity in age-matched samples, and that developmental growth of MG is highly consistent (**Fig. 2C**; CoV 24 hpf 18.72%, 48 hpf 3.13%, 60 hpf 5.25%, 72 hpf 6.52%; p<0.0001).

**Figure 2.**
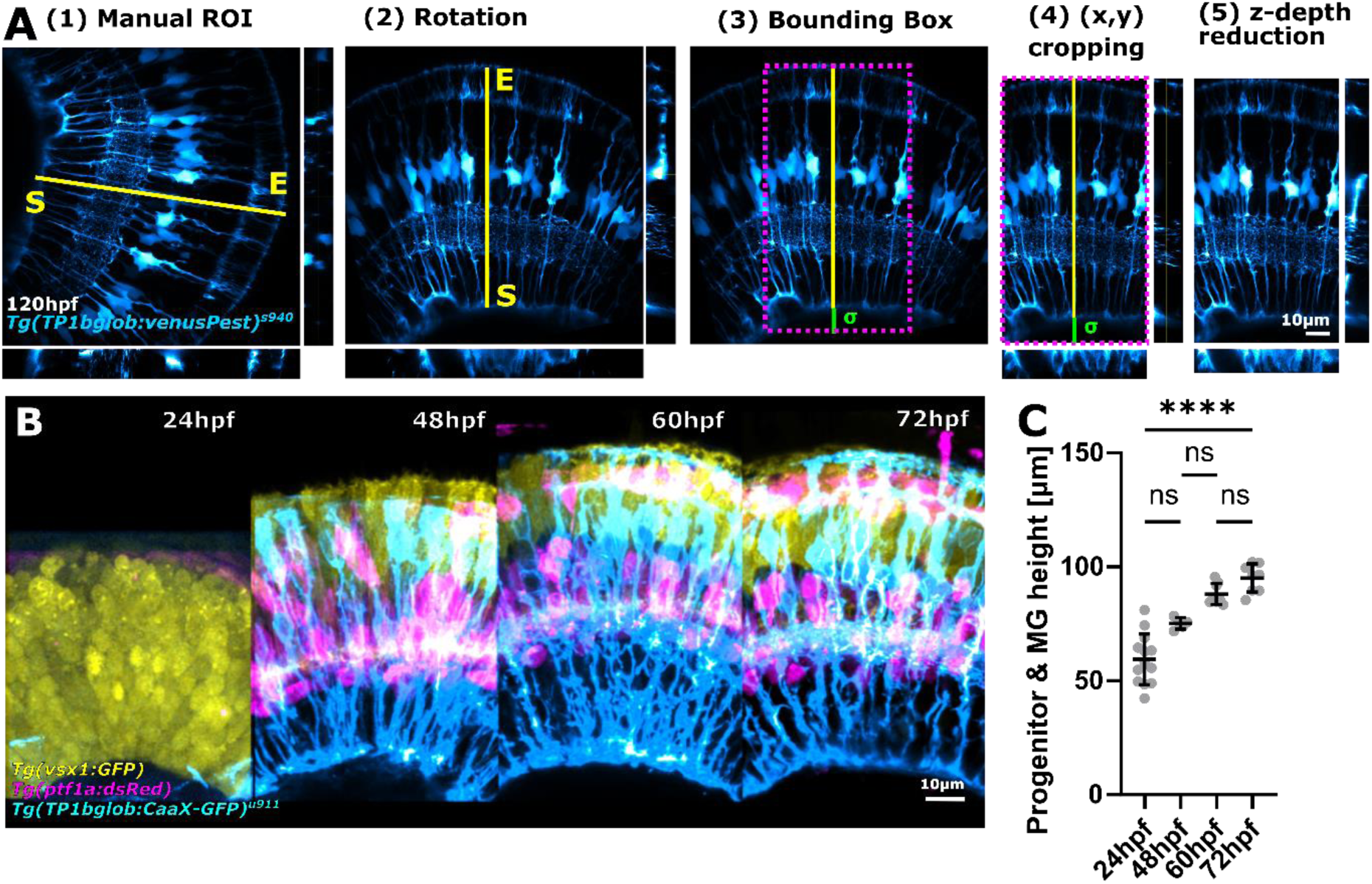
Image standardization is required for image comparability. **(A)** The subregionTool allows for semi-automatic subregion selection by (i) manual line-ROI selection, which is then used to rotate the image (ii), create a bounding box (iii), crop the image using this bounding box (iv), and crop the image in z (v) (*sigma = user-defined basal extension to allow for blood vessel inclusion*). **(B)** Applying the subregionTool to images from different transgenic reporter lines, shows that they can be made comparable, as exemplified by computationally overlapping them. **(C)** Measurement of progenitor/MG height shows a high similarity in age-matched samples, and that developmental growth of MG is highly reproducible between samples (CoV 24hpf 18.72%, 48hpf 3.13%, 60hpf 5.25%, 72hpf 6.52%; p<0.0001; 24hpf n=13 embryos, 48hpf n=5 embryos, 60hpf n=8 embryos, 72 hpf n=8 embryos; N=2 experimental repeats; Kruskal-Wallis test; mean ± s.d.).

Reproducible 3D ROIs with minimal user input allowed for the establishment of image comparability objectively, making sample and group assessments possible. Together, this facilitates the comparison of MG morphology between images and datasets.

### The image enhancement workflow depends on the acquisition type

When performing fluorescence microscopy, typically the image does not directly reproduce the object of interest due to artefacts as well as the system impulse function or convolution of light, called point spread function (PSF) [33]. To restore the object properties before data processing and object measurements, typically a deconvolution step is performed. Often deconvolution is done with external software (such as Huygens from Scientific Volume Imaging), but this can also be achieved with Fiji using external Plugins. We next examined PSF deconvolution in Fiji and used this information to design the ***deconvolutionTool*** for GliaMorph. For data acquired in confocal mode (**Fig. S3A**), theoretical PSFs were established based on transgenic reporter line-specific fluorophores. Also, the following PSF deconvolution algorithms were examined, namely Regularized Inverse Filter (**Fig. S3B**), Landweber (**Fig. S3C**), Fast Iterative Shrinkage Thresholding (**Fig. S3D**), Bounded-Variable Least Squares (**Fig. S3E**), and Richardson-Lucy (RL; **Fig. S3F**), finding RL to be the most applicable to our data.

Next, the above deconvolution was implemented into the *deconvolutionTool* which allows single-/multi-channel input, selection of fluorophore wavelengths, different objective numerical aperture (NA), and theoretical or experimental PSF file input. The *deconvolutionTool* can be applied to any data acquired on traditional confocal microscopes.

As Zeiss AiryScan microscopy allows for sub-diffraction resolution, allowing for increased detail and increased CNR, we next acquired data with this imaging paradigm. As expected, this showed 3D deconvolution to outperform 2D deconvolution, resulting in reduced background (white arrowheads), and high deconvolution to result in increased structured noise and grains (black arrowheads; **Fig. S4**).

Together, this emphasizes that not only the imaging (*i.e.,* confocal vs AiryScan) but also the selected type and parameters of deconvolution can influence data quality for subsequent data analysis.

### Müller glia apicobasal texture can be visualized using 1D-vectors

We examined whether it was possible to discern different MG zones given the fact that MG show a stereotypic apicobasal pattern. This was since MG patterns can computationally be seen as a differential distribution of geometries and intensity (**Fig. 3A**). However, as the examined data are 3-dimensional (3D), we explored whether dimensionality reduction from 3D (x,y,z) data to a 1D vector would allow for data exploration. Thus, we reduced data using the ***zonationTool*** (**Fig. 3B**). This tool reduces data first in the z-axis, transforms them by 90°, and then performs another dimensionality reduction. This allows assessment of the intensity distributions or “zonation” from apical-to-basal across the retina. As the approach is independent of input data, it can be applied to different labelling approaches of MG as well as other cell types, such as retinal neurons (**Fig. 3C-F**).

**Figure 3.**
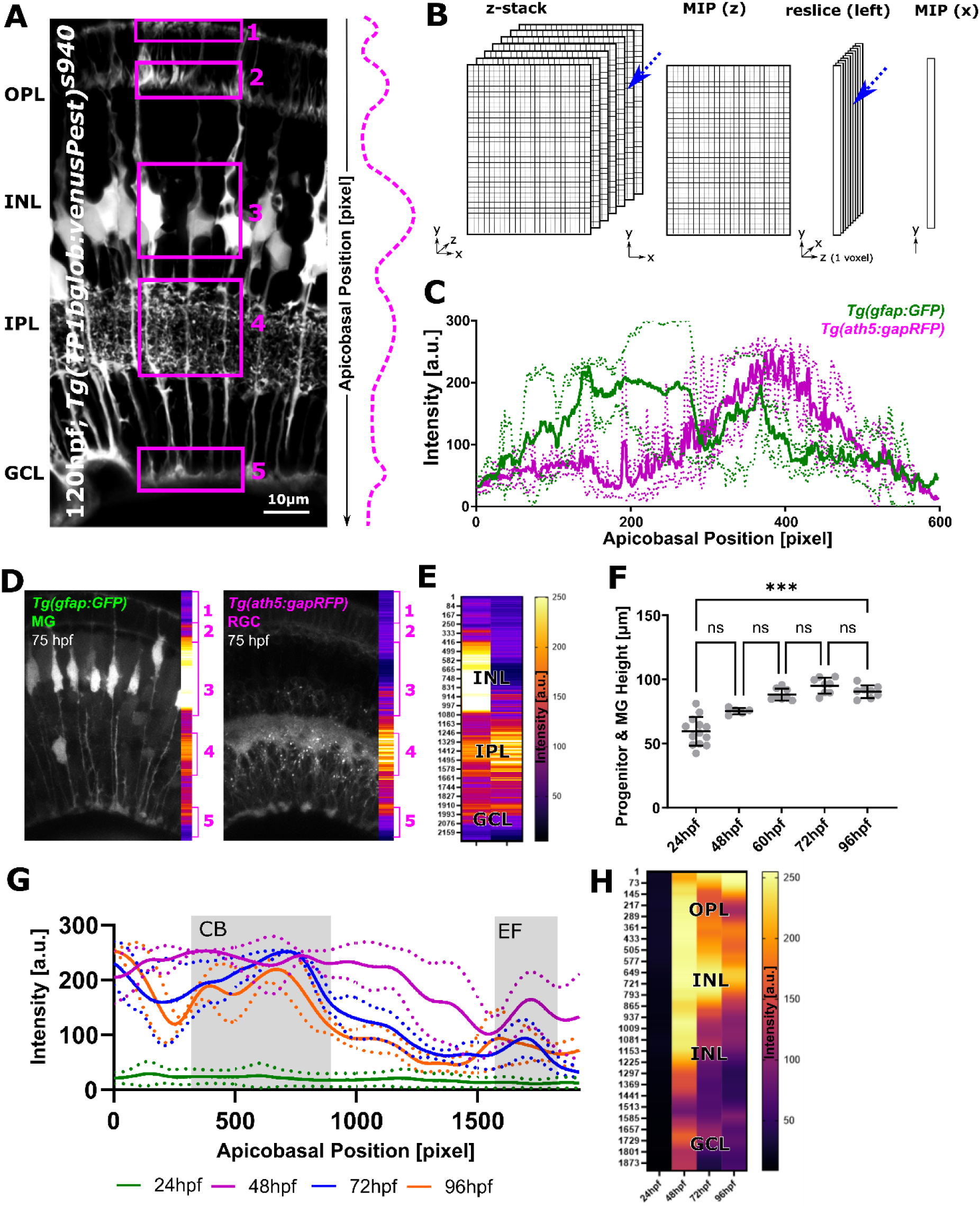
The ZonationTool enables data reduction to 1D-vectors for apicobasal texture analysis. **(A)** MG have subcellular morphological specialisations from the apical (top) to basal (bottom) position in the retina. These subregions (1-5), highlighted in boxes, were expected to be discernible by distinctive intensity profiles (such as high levels for nuclei and lower levels for the IPL). **(B)** Diagram of the workflow: 3D image stacks were reduced to 2D images, transformed to 1D+z, and again reduced, thus resulting in a one-voxel-wise representation of MG data. **(C)** Apicobasal intensity plot of a double- transgenic at 75hpf, shows that the subregionTool applies to MG as well as neurons. Application to double-transgenics moreover allows understanding of cell relationships (n=3). **(D)** Representative images used for plots in F, showing differences between transgenics. **(E)** Heatmap representation for apicobasal texture analysis of D. **(F)** Retina height measurements from 24-to-96 hpf showing a statistically significant increase over time (p=0.0006; not significant (ns) p>0.999; 24hpf n=13 embryos, 48hpf n=5 embryos, 60hpf n=8 embryos, 72hpf n=8 embryos, 96hpf n=8 embryos; N=2 experimental repeats; Kruskal-Wallis test; mean ± s.d.). **(G)** Intensity profiles from 24- to-96hpf produced with the zonationTool (solid line depicting mean values; image size normalized to 1900 for comparability). **(H)** Heatmap representation for apicobasal texture analysis of H (mean).

We next examined whether MG at different developmental stages are comparable in size and how this changed over time. There was a significant increase of MG height from 24-to-96 hpf (p=0.0006, Kruskal-Wallis test; **Fig. 3F**). When analysing the coefficient of variation (CoV), variation was low, suggesting comparability between samples. Briefly, highest CoV was observed at 24 hpf 18.72%, 48 hpf 3.31%, 60 hpf 5.25%, 72 hpf 6.52%, and 96 hpf 5.46%. Analysing MG texture from 24-to-96 hpf, with normalization to 1920 px to allow direct visual comparison, showed that endfeet are identifiable from 48 hpf onwards. A clear discrimination between MG cell bodies and IPL is possible from 72 hpf onwards (**Fig. 3G,H**). Together, dimensionality reduction of 3D images using the *zonationTool* is a user-friendly way to visualize texture and subregional zones of MG as 1D-vectors. Due to its independence of input data, it applies to any samples, if image comparability is established.

### GliaMorph allows 3D feature extraction and quantification of MG

*Tg(TP1bglob:VenusPest)^s940^* is an established transgenic reporter line to visualize MG, here used this to develop and test the GliaMorph toolkit to quantify 3D cell features. To examine whether MG features elaborated with maturity, we analyzed data at 72 hpf and 120 hpf. After using the *SubregionTool* for data comparability, the *ZonationTool* was used to plot the apicobasal texture of MGs. This showed growth of MG in size and downward migration of nuclei (**Fig. 4A**; cell subdomain 3). To allow for the assessment of pattern vs intensity-level differences, the zonationTool allows for normalization (**Fig. 4A-C**). Thus, one can directly assess intensity levels of treatments against each other. For example, one could examine this further using histograms for level and frequency; use area under the curve measurements for peaks and distributions; use correlations for more local analysis.

**Figure 4.**
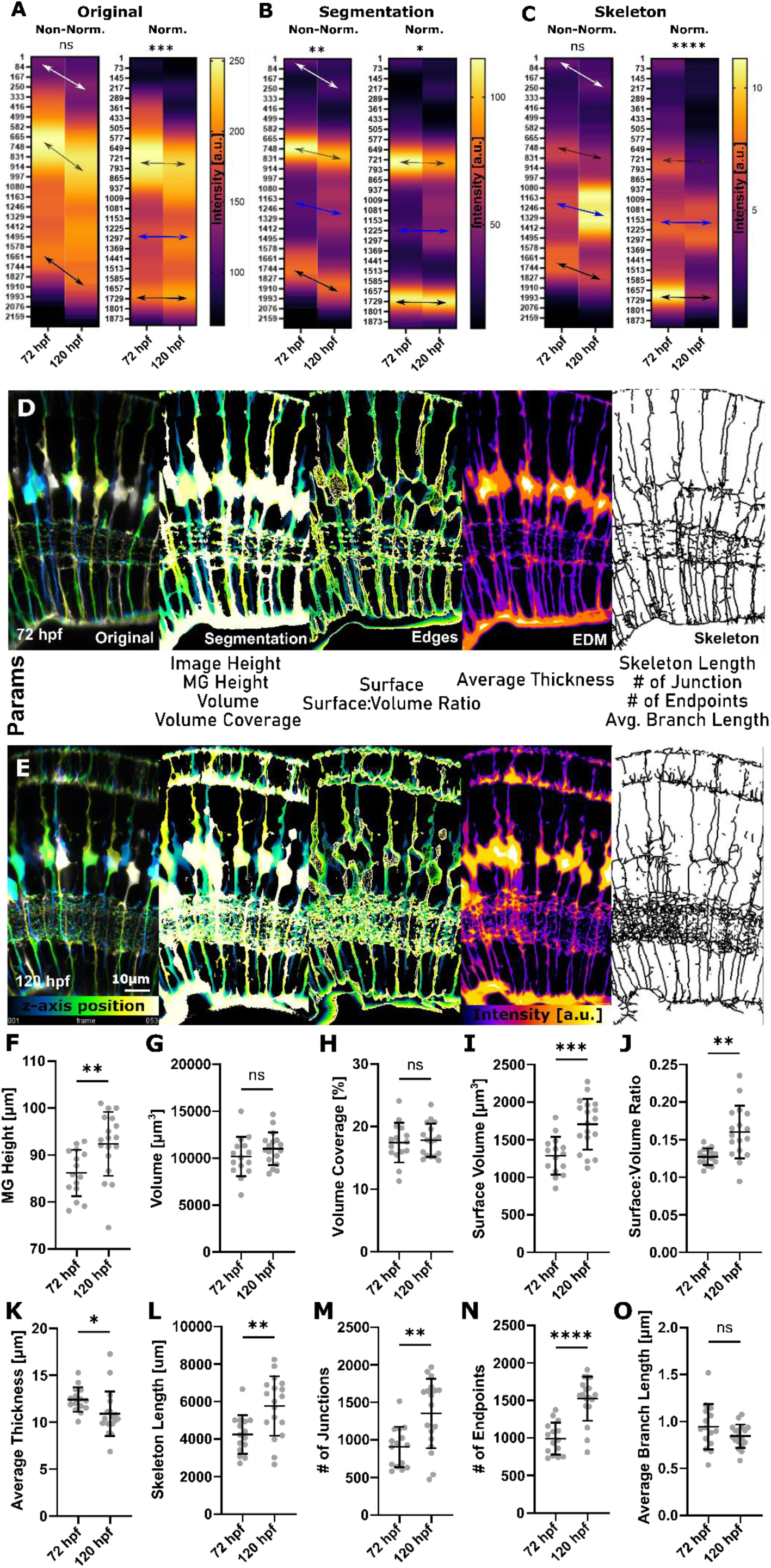
GliaMorph allows 3D feature extraction and quantification of MG. **(A)** Apicobasal texture plot of original images showing the maturation of MG as indicated by changes in subregion 1/2 (white arrow), cell bodies (grey arrow), and endfeet (p=0.0862; Mann-Whitney U test; mean). Normalization refers to image length, *i.e.* both images were adjusted to the same length. **(B)** Apicobasal texture plot of segmented images shows similar changes as observed when plotting original data, but also indicates IPL maturation (blue arrow) (p=0.0052; Mann-Whitney U test; mean). **(C)** Apicobasal texture plot of skeletonized images, again shows that MG the network elaborates over time (p=0.3402; Mann-Whitney U test; mean). **(D,E)** Workflow overview to extract MG features on a 3D global image-level, including depth-coded (DC) original images, DC segmentation, thickness (Euclidean distance map; higher intensity represents thicker regions), skeleton (MIP dilated for representation), and DC surface at 72 hpf and 120 hpf. **(F,O)** MG morphology significantly matures between 72hpf and 120hpf. **(F)** MG height was significantly increased from 72-to-120 hpf (p=0.0061; two-tailed unpaired t-test; mean ± s.d.). **(G)** Volume was not significantly changed (p=0.2314; two-tailed unpaired t-test; mean ± s.d.). **(H)** Volume coverage was not significantly changed from (p>0.9999; two-tailed unpaired t-test; mean ± s.d.). **(I)** Surface volume was significantly increased (p=0.0004; two-tailed unpaired t-test; mean ± s.d.). **(J)** Surface-to-volume ratio was statistically significantly increased (p=0.0015; two-tailed unpaired t-test; mean ± s.d.). **(K)** Average thickness was significantly decreased (p=0.0366; two-tailed unpaired t-test; mean ± s.d.). **(L)** Skeleton length was statistically significantly increased (p=0.0033; two-tailed unpaired t-test; mean ± s.d.). **(M)** Number of junctions was significantly increased (p=0.0025; two-tailed unpaired t-test; mean ± s.d.). **(N)** Number of endpoints was significantly increased (p=<0.0001; two-tailed unpaired t-test; mean ± s.d.). **(O)** Average branch length was not significantly changed (p=0.1409; 72 hpf n=15, 120 hpf n=18; N=2 experimental repeats; two-tailed unpaired t-test; mean ± s.d.).

To extract MG in the images, we established the ***segmentationTool*** that uses bleach correction, 8-bit conversion, 3D Median filtering, and Otsu-based thresholding to produce binary/segmented images. We then applied the ***quantificationTool***, which extracts the following global image-level features from the segmented image: **(i)** image height: length of y-axis, **(ii)** MG volume: voxels classified as MG after segmentation, **(iii)** density: ratio of total image voxels divided by MG volume voxels (*i.e.* given as a fraction of 1). After surface extraction using Canny edge detection, **(iv)** surface area is quantified. Using 3D-thinning, the skeleton/centreline was extracted to quantify **(v)** network length, **(vi)** number of branching points, **(vii)** number of end points, and **(viii)** average branch length. Lastly, combining the skeleton with a 3D Euclidean Distance Map**, (ix)** the average thickness was analysed.

Applying the *zonationTool* to the segmented data showed again an increase in size and MG cell bodies that were positioned more basally at 120 hpf. Also, clear bands of apical MG zones (1 and 2) and endfeet are seen (**Fig. 4B**; subdomain 5). When plotting the automatically skeletonized images, elaborations of MG from 72-to-120 hpf were pronounced in the IPL (**Fig. 4C**; subdomain 4). We quantified MG features with the *quantificationTool* at 72 hpf and 120 hpf (**Fig. 4D,E**). Quantification of MG height showed a significant increase from 72-to-120 hpf (p=0.0061, **Fig. 4F**). Surprisingly, neither MG volume (p=0.2314, **Fig. 4G**) nor percentage volume coverage were significantly different from 72-to-120 hpf (p>0.9999, **Fig. 4H**). However, MG surface volume (p=0.0004, **Fig. 4I**) and surface-to-volume ratio were significantly increased from 72-to-120 hpf (p=0.0015, **Fig. 4J**), which suggested that shape complexity increased over time. The average thickness was significantly decreased from 72-to- 120 hpf (p=0.0366, **Fig. 4K**), which was thought to be due to an increase in the number of thinner protrusions over time. As expected, skeleton length (p=0.0033, **Fig. 4L**), number of junctions (p=0.0025, **Fig. 4M**), and number of endpoints were statistically significantly increased from 72-to-120 hpf (p<0.0001, **Fig. 4N**), while average branch length was not statistically significantly altered (p=0.1409, **Fig. 4O**).

Together, this shows that GliaMorph is suitable to assess MG morphology in complete transgenic retinas in 3D and allows extraction of biologically meaningful information.

### Visualization of MG membranes supersedes detail visualized with cytosol reporters

As GliaMorph analysis is based on object intensity and distribution, we next compared cytosolic or membrane markers for 3D MG morphological analysis. Consequently, a membrane marker construct was injected into a cytosolic transgenic to achieve mosaic expression to visualize individual cells (72 hpf; **Fig. S6A-C**). Visually, the membrane-marker delineated more detail than the cytosol-marker. This is exemplified in regions such as MG protrusions in the IPL (white arrowheads) or MG honeycombing or anisotropic scaffolding [34] in the OLM (unfilled arrowhead). The segmentation approach delivered satisfying outcomes with the membrane-marker, but not the cytosol-marker (**Fig. S6D-F**); moreover, with the membrane-marker cell connectivity and IPL protrusion details were extracted. This was also reflected in the 3D skeleton, which showed more detail with the membrane-marker (**Fig. S6G-I**) and resolved cell domains to a satisfactory level (see **Fig. 1C**).

Hence, the “ideal” situation was found to be individual MG clones labelled with a membrane marker for detail. However, as producing clones can be laborious and not uniform across different injected animals, we next studied a global transgenic with membrane-labelled MG *Tg(TP1bglob:eGFP-CAAX) ^u911^*.

### MG development is defined by apicobasal elaboration and refinement

As membrane labels outperformed cytosolic MG cell labelling in clones, we generated the stable transgenic line *Tg(TP1bglob:eGFP-CAAX)^u911^.* Using this, we analysed MG development in a shorter time-frame from 60-to-96 hpf (**Fig. 5A-C**). This revealed statistically significant increases in MG height (p<0.0001; **Fig. 5D**), thickness (p=0.0466; **Fig. 5I**), and average branch length (p=0.0018; **Fig. 5M**), but none of the other measured parameters.

**Figure 5.**
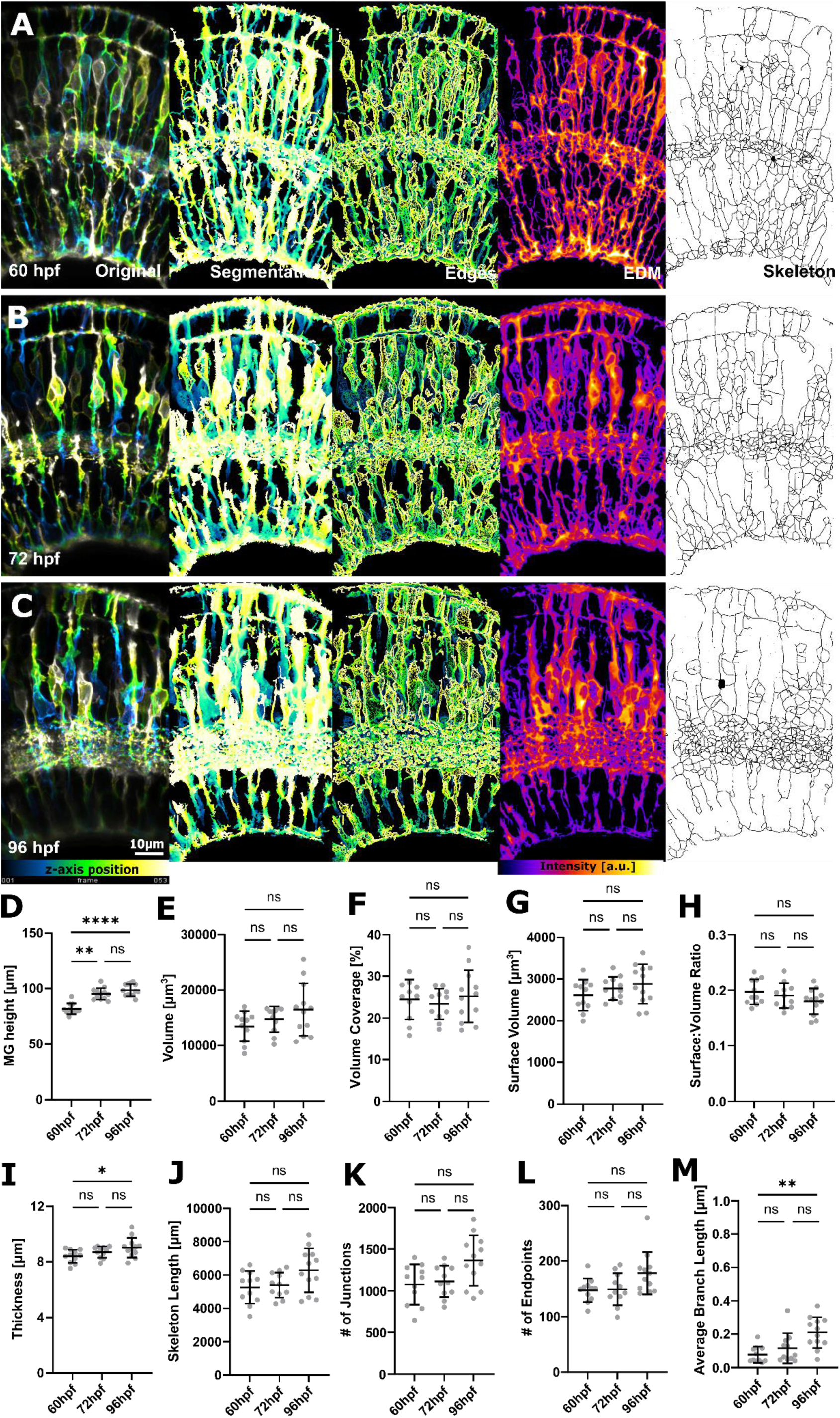
MG global feature analysis in a stable membrane-visualizing transgenic. **(A-C)** Micrographs of original and processed data at 60, 72, and 96 hpf, respectively. **(D-M)** All parameters, except MG height, did not significantly change during the studied time frame. **(D)** MG height was statistically significant increased from 60-96hpf (p<0.0001; Kruskal-Wallis test; mean ± s.d.). **(E)** Volume was not statistically significantly changed (p=0.2197; Kruskal-Wallis test; mean ± s.d.). **(F)** Volume coverage was not statistically significantly changed (p=0.7728; Kruskal-Wallis test; mean ± s.d.). **(G)** Surface volume was not statistically significantly changed (p=0.3036; Kruskal-Wallis test; mean ± s.d.). **(H)** Surface-to-volume ratio was not statistically significantly changed (p=0.3570: Kruskal-Wallis test; mean ± s.d.). **(I)** Thickness was statistically significant increased from 60-96hpf (p=0.0466; Kruskal-Wallis test; mean ± s.d.). **(J)** Skeleton length was not statistically significantly changed (p=0.1095; Kruskal-Wallis test; mean ± s.d.). **(K)** Number of junctions was not statistically significantly changed (p=0.0741; Kruskal-Wallis test; mean ± s.d.). **(L)** Number of endpoints was not statistically significantly changed (p=0.0690; Kruskal-Wallis test; mean ± s.d.). **(M)** Average branch length was statistically significant increased from 60-96hpf (p=0.0018; 60hpf n=11, 72hpf n=12, 96hpf n=13; N=2 experimental repeats; Kruskal-Wallis test; mean ± s.d.).

This led us to examine our data in a local fashion using apicobasal distributions. This revealed a significant difference in intensities from 60-to-96 hpf for original (p<0.0001), segmented (p<0.0001), and skeletonized images (p<0.0001; **Fig. 6A**). We observed a downward migration of nuclei, increased MG height, and increased overall complexity (as indicated by skeleton distributions), particularly in the IPL. We then examined the alignment of structures in the image (*i.e.,* horizontal vs vertical) that can be described as image order [35] (**Fig. 6B,C**), which showed a significant difference from 60-to-96 hpf (p=0.0049; **Fig. 6D**). These data suggests that even though features might not change enough to be extracted globally (**Fig. 5**), local features are elaborated and refined over time. As these measurements are based on the population-level (image-level or global) analysis, we next sought to study cell heterogeneity and whether measurements of individual cells represent collective measurements of cell populations. Thus, we visualized and analysed data from individual clones from 60-to-96 hpf (**Fig. 7A**). In individual MGs, we saw changes in retina height, surface-to-volume ratio, skeleton length, and endpoints (**Fig. 7B,E,H**) just as we observed at the population level (**Fig. 4D,H,I,L**). Other parameters such as volume, surface, and number of junctions were increased in single-cell level measurements (**Fig. 7C,D,G**) and branch length was not altered on the single-cell level (**Fig. 7I**).

**Figure 6.**
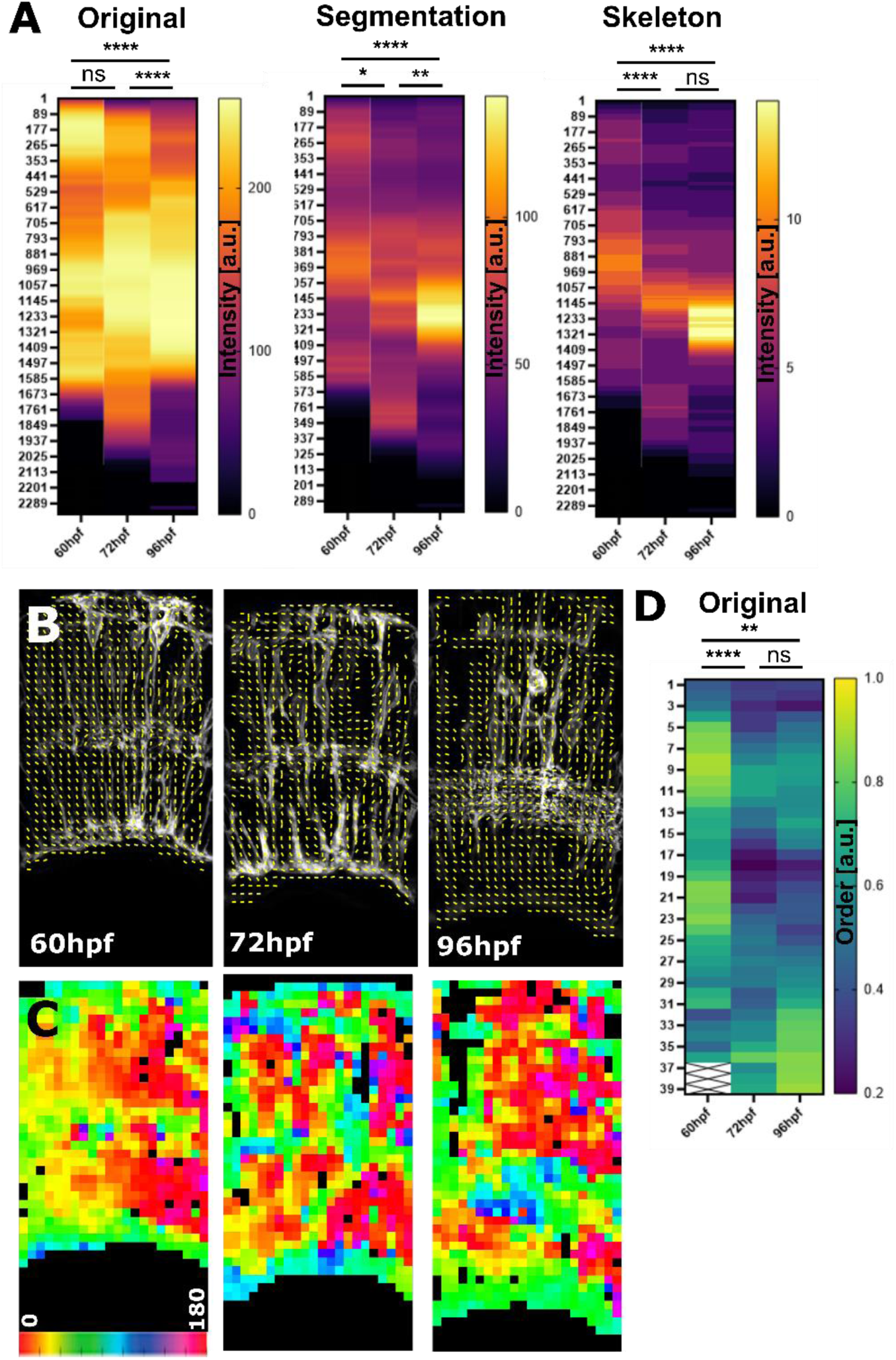
MG higher-order analysis in a stable membrane-visualizing transgenic. **(A)** Apicobasal intensity plotting showed a statistically significant difference from 60- 96 hpf in original (p<0.0001), segmented (p<0.0001), and skeletonized images (p<0.0001; 60hpf n=11, 72hpf n=12, 96hpf n=13; N=2 experimental repeats; Kruskal-Wallis test; mean). This suggested that subcellular features matured over time. **(B-D)** Using measurements of orientation, showed that the subcellular organization changed from a more vertical (1-yellow) to a more horizontal (0.2 – blue) alignment. (B – vectors; C - colourized representation of vector angles; D - image order was statistically significantly different from 60-96hpf (p=0.0049; data from 2 experimental repeats; Kruskal-Wallis test)).

**Figure 7.**
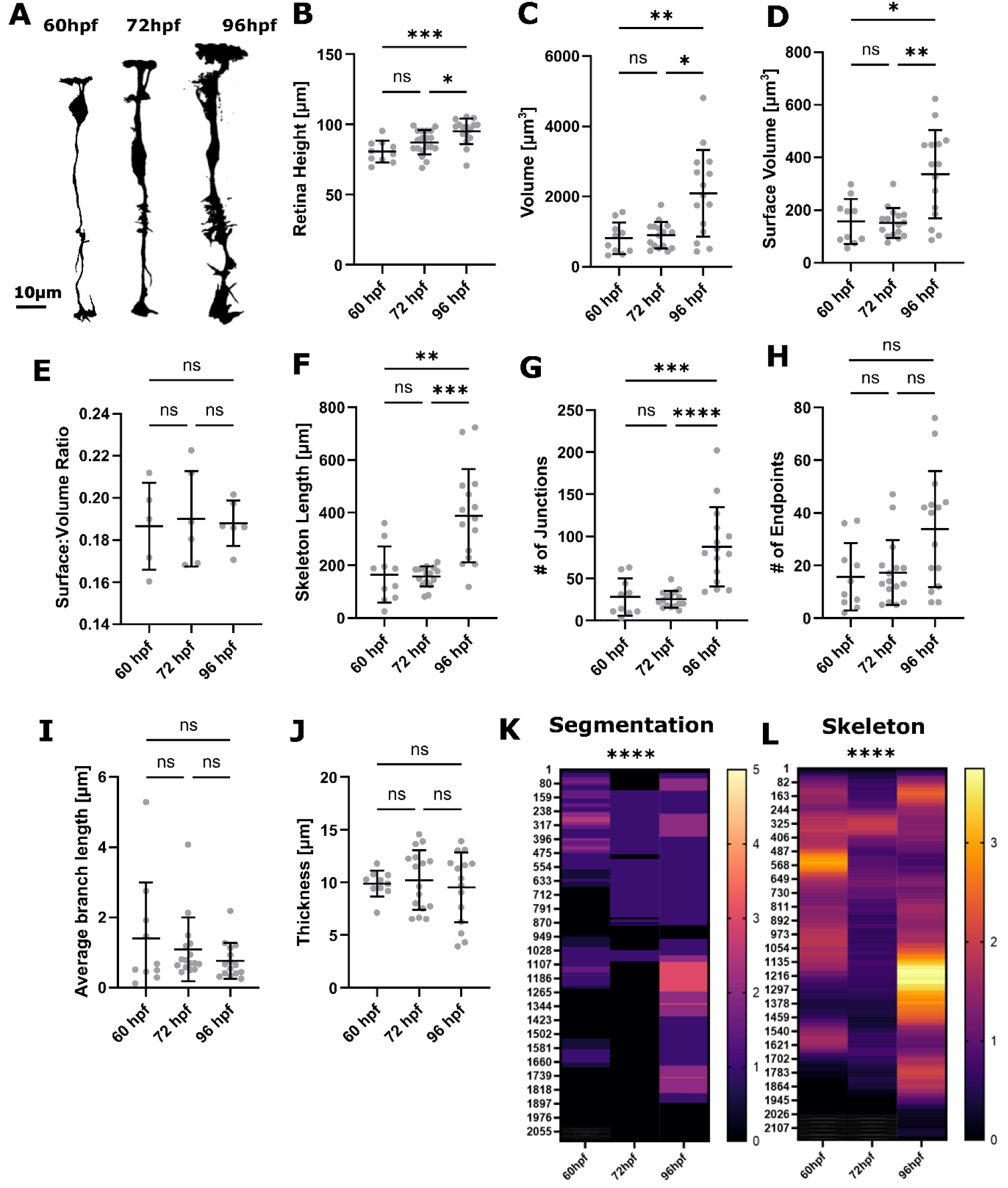
Analysis of glial development using single-cell measurements. **(A)** Segmentation MIPs of clones at 60hpf, 72hpf, and 96hpf clones (representative images extracted from 3D stacks). **(B)** MG height did significantly increase from 60- to-96hpf (p=0.0006; Kruskal-Wallis test; mean ± s.d.). **(C)** Volume did significantly increase from 60-to-96hpf (p=0.0035; Kruskal-Wallis test; mean ± s.d.). **(D)** Surface volume did significantly increase from 60-to-96hpf (p=0.0029; Kruskal-Wallis test; mean ± s.d.). **(E)** Surface-to-volume ratio did not significantly change from 60-to-96hpf (p=0.9947; mean ± s.d.). **(F)** Skeleton length did significantly increase from 60-to- 96hpf (p<0.0001; mean ± s.d.). **(G)** Number of junctions did significantly increase from 60-to-96hpf (p<0.0001; mean ± s.d.). **(H)** Number of endpoints did not significantly alter from 60-to-96hpf (p=0.0400; mean ± s.d.). **(I)** Average branch length did not significantly alter from 60-to-96hpf (p=0.3320; mean ± s.d.). **(J)** Thickness did not significantly alter from 60-to-96hpf (p=0.8241; 60hpf n=10 cells, 72hpf n=19 cells, 96hpf n=16 cells; N=3 experimental repeats; Kruskal-Wallis test; mean ± s.d.). **(K,L)** Apicobasal intensity plotting showed a statistically significant difference from 60-96hpf in segmented (p<0.0001), and skeletonized images (p<0.0001).

Together, our data show that single cell and global image-level measurements are in good agreement, but that batch effects might impact measurement outcomes. For example, single-cell thickness measurements showed high variability, which was not observed for global image-level measurements. Conversely, using single-cell measurements allows for a closer examination of cellular heterogeneity. While precise measurements can be derived from single-cell analysis, the sampling problem they introduce becomes important. This highlights that global and single-cell analysis might answer different biological questions.

### Loss of *cadherin2* leads to altered MG apicobasal feature distribution

We next tested whether GliaMorph would allow us to describe disease-associated glia changes. Hence, *cadherin2* (*cdh2* or *N-cadherin*) was knocked out using CRISPR/Cas9 technology [36]. We chose *cdh2* as it is a cell surface adhesion molecule shown to play a role in basal migration of retinal progenitors and MG formation upon injury in adult zebrafish retinas [23], [37]. This suggested that cdh*2* may play a role in the establishment of MG morphology. When quantifying MG features on the global level, we saw no significant difference in 3D measurements (**Fig. 8C-L**). However, analysing the apicobasal texture in the same images, MG were found to be significantly different in the original (p<0.0001), segmented (p<0.0001), and skeletonized (**Fig. 8M**; p=0.0072) data. To further understand this, we performed directionality analysis based on Fourier Transformation as previously described [35] (**Fig. 8N**). This revealed a significant difference in the vertical-to-horizontal cell alignment (p=0.0221; **Fig. 8O**), particularly at the apex of MGs and within the IPL. This suggests that loss of *cdh2* did not lead to significant overall changes in MG size or volume, but that the organization of cell elaboration orientation is impaired. It is important to note that the examined crispants were mosaic mutants and unlikely to have a highly penetrant phenotype, which could make it challenging to identify subtle MG shape changes. To examine whether this is mirrored in single-cell analysis, we performed the same analysis using single MG clones generated by injection of DNA constructs with gRNAs (**Fig. S7**). While MG height was found to be increased in *cdh2* crispants (p=0.0049; **Fig. S8C**), the other parameters were not changed - which was in agreement with global image-level measurements (**Fig. S7C-J**). Apicobasal measurements again showed a significant difference (**Fig. S7K-L**; both, p<0.0001).

**Figure 8.**
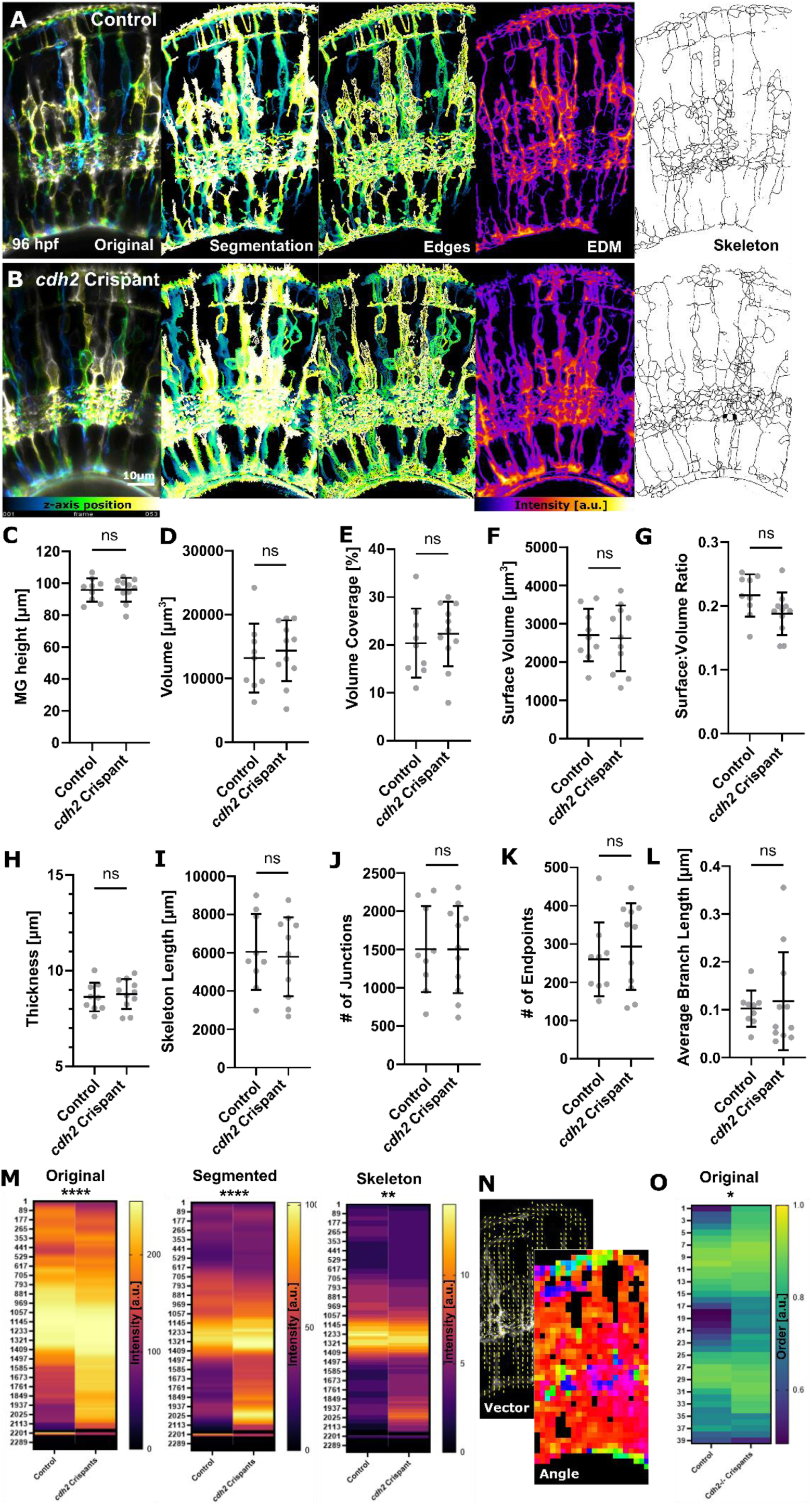
Loss of *cadherin 2* alters apicobasal features. **(A, B)** Maximum intensity projections (MIPs) of controls and *cadherin 2* crispants, visualizing MG with the transgenic reporter line *Tg(TP1bglob:eGFP-CAAX) ^u911^.* **(C-L)** Analysis of global image-level features showed no significant difference upon *cadherin 2* loss. **(C)** MG height was not significantly changed upon loss of cadherin 2 (p=0.9519; control n=9, *cdh2^-/-^* n=11; N=2 experimental repeats; unpaired Students t-test; mean ± s.d.). **(D)** MG volume was not significantly changed upon loss of cadherin 2 (p=0.6211; control n=9, *cdh2^-/-^* n=11; N=2 experimental repeats; unpaired Students t- test; mean ± s.d.). **(E)** Volume coverage was not significantly changed upon loss of cadherin 2 (p=0.5461; unpaired Students t-test; mean ± s.d.). **(F)** MG surface volume was not significantly changed upon loss of cadherin 2 (p=0.8175; unpaired Students t-test; mean ± s.d.). **(G)** MG surface-to-volume ratio was not significantly changed upon loss of cadherin 2 (p=0.0725; unpaired Students t-test; mean ± s.d.). **(H)** Average thickness was not significantly changed upon loss of cadherin 2 (p=0.6422; unpaired Students t-test; mean ± s.d.). **(I)** Skeleton length was not significantly changed upon loss of cadherin 2 (p=0.7827; unpaired Students t-test; mean ± s.d.). **(J)** Number of junctions was not significantly changed upon loss of cadherin 2 (p=0.9837; unpaired Students t-test; mean ± s.d.). **(K)** Number of endpoints was not significantly changed upon loss of cadherin 2 (p=0.4922 unpaired Students t-test; mean ± s.d.). **(L)** Average branch length was not significantly changed upon loss of cadherin 2 (p=0.6845; unpaired Students t-test; mean ± s.d.). **(M)** Apicobasal texture was significantly changed when looking at original (p<0.0001; Mann-Whitney U test), segmented (p<0.0001; Mann-Whitney U test), and skeletonized (p=0.0072; Mann-Whitney U test; mean) data. This suggested that subcellular components were changed upon loss of *cadherin 2*. **(N, O)** Using image orientation measurements, showed that feature alignment was significantly altered upon *cadherin 2* loss – this can be particularly seen in the OLM and IPL (p=0.0221; control n=9, *cdh2^-/-^* n=11; N=2 experimental repeats; Mann Whitney U test; mean).

Together, we show that apicobasal feature analysis is sufficient to detect cell shape alterations upon loss of *cdh2* on a global as well as single-cell level. Using orientation assessment in addition to apicobasal feature analysis shows that loss of *cdh2* does not only lead to overall shape changes but also affects cell alignment. These data suggest that GliaMorph could be used to screen for novel molecules involved in glia shape.

### Apicobasal feature analysis provides insights into mouse glaucoma models

So far, we applied GliaMorph to transgenic reporter lines labelling MG in the zebrafish retina. However, many studies where MG morphology is of interest (e.g., in diseased tissues) may use other models, such as mice, and do not always have a cell-specific fluorescent transgenic reporter available. Thus, we tested GliaMorph on the retina from another species (*i.e.*, mice) where MG are visualized with antibody staining. To assess whether biologically relevant data could be extracted, we collected retinas from CD1 controls and DBA/2J mice, which develop glaucoma-like phenotypes and exhibit gliosis [24]. We used the Rlbp1/Cralbp antibody to label the entire MG cell and GFAP antibody, which is a commonly used marker for gliosis and an indicator of pathology in retinal degenerative diseases (**Fig. 9A,B**). When analysing apicobasal distributions, we found changes in Rlbp1, suggesting that cell morphology and subcellular arrangements are changed in this glaucoma model (**Fig. 9C**; original p<0.0001, segmentation p<0.0001, skeleton p<0.0001). Additionally, we found GFAP to be upregulated and distributed to a more apical area in glaucoma mice in comparison to controls (**Fig. 9D**; original p<0.0001; segmentation p<0.0001; skeleton p<0.0001). Volume, branching, and size quantifications using the GliaMorph suite showed no statistically significant difference (**Fig. S8)**.

**Figure 9.**
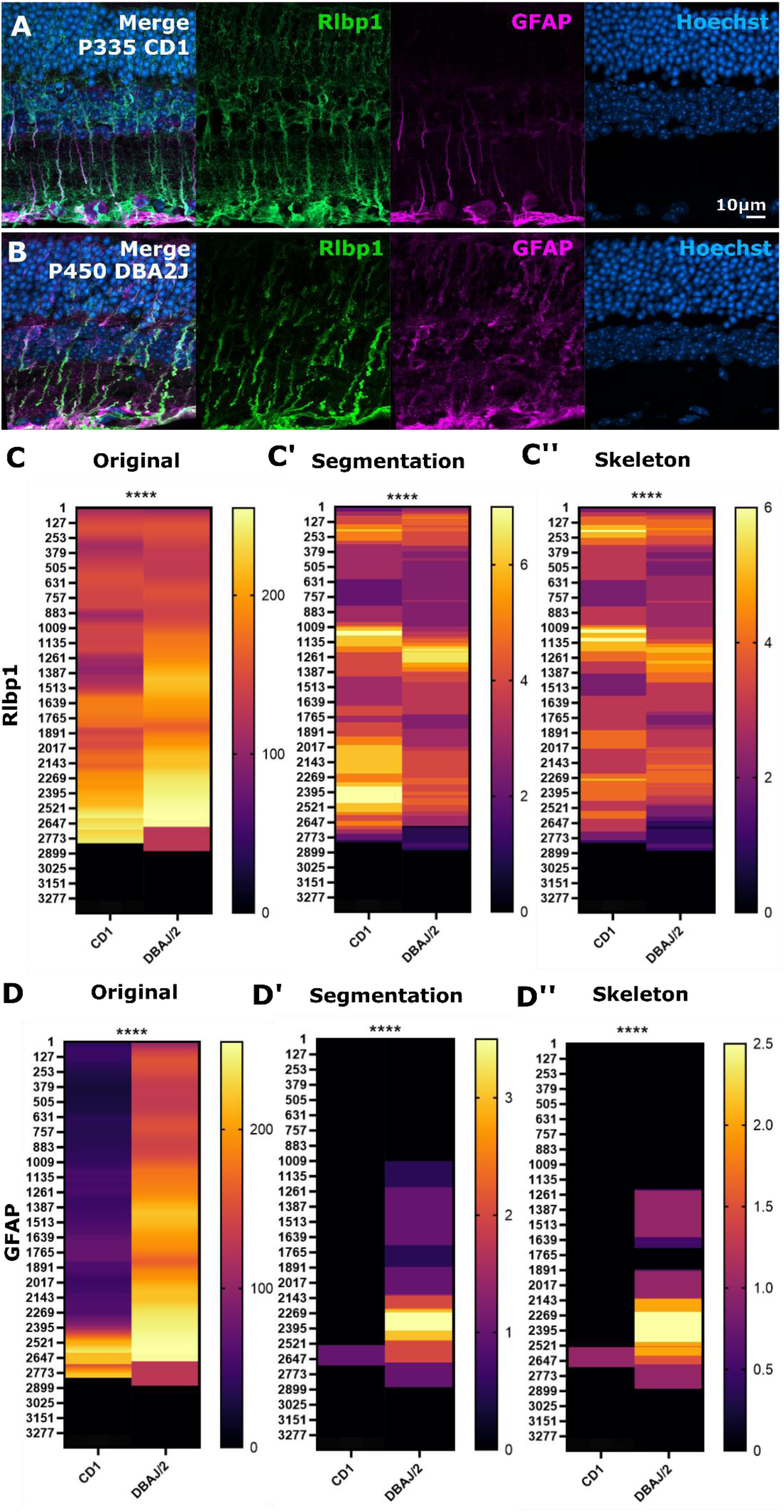
Apicobasal texture analysis can be used to study mouse glaucoma disease models. **(A, B)** Micrographs of controls and spontaneously glaucomatous DBA/2/J mice. **(C)** Using Rlbpj1 staining to visualize the complete MG structure, shows structural changes in glaucoma when analysed using apicobasal texture analysis (original p<0.0001; segmentation p<0.0001; skeleton p<0.0001; Mann-Whitney U test; n=7 stacks from 3 mice each). **(D)** Using GFAP staining to visualize reactive MG with cytoskeletal changes, shows structural changes in glaucoma when analysed using apicobasal texture analysis (original p<0.0001; segmentation p<0.0001; skeleton p<0.0001; Mann-Whitney U test; n=7 stacks from 3 mice each).

Together, this shows that GliaMorph can be used with images from retinas of different species and antibody staining to observe and quantify MG phenotypes. Specifically, the GliaMorph texture analysis provides a crucial method to identify MG phenotypes quickly and robustly in pathology samples stained with antibodies.

### Workflow integration

The GliaMorph toolkit allows the workflow to be individualized for experimental needs based on its modular construction (**Fig. 10**). Batch processing allows the selected tools to be run on whole experimental folders, increasing throughput and automation. Implementation in the Fiji framework allows cross-platform and licence-free applicability. Implementing codes as Macros with GUIs allows direct use and alteration of code, even by users without any coding experience. This is supported by supplying example data as well as a step-by-step user guide.

**Figure 10.**
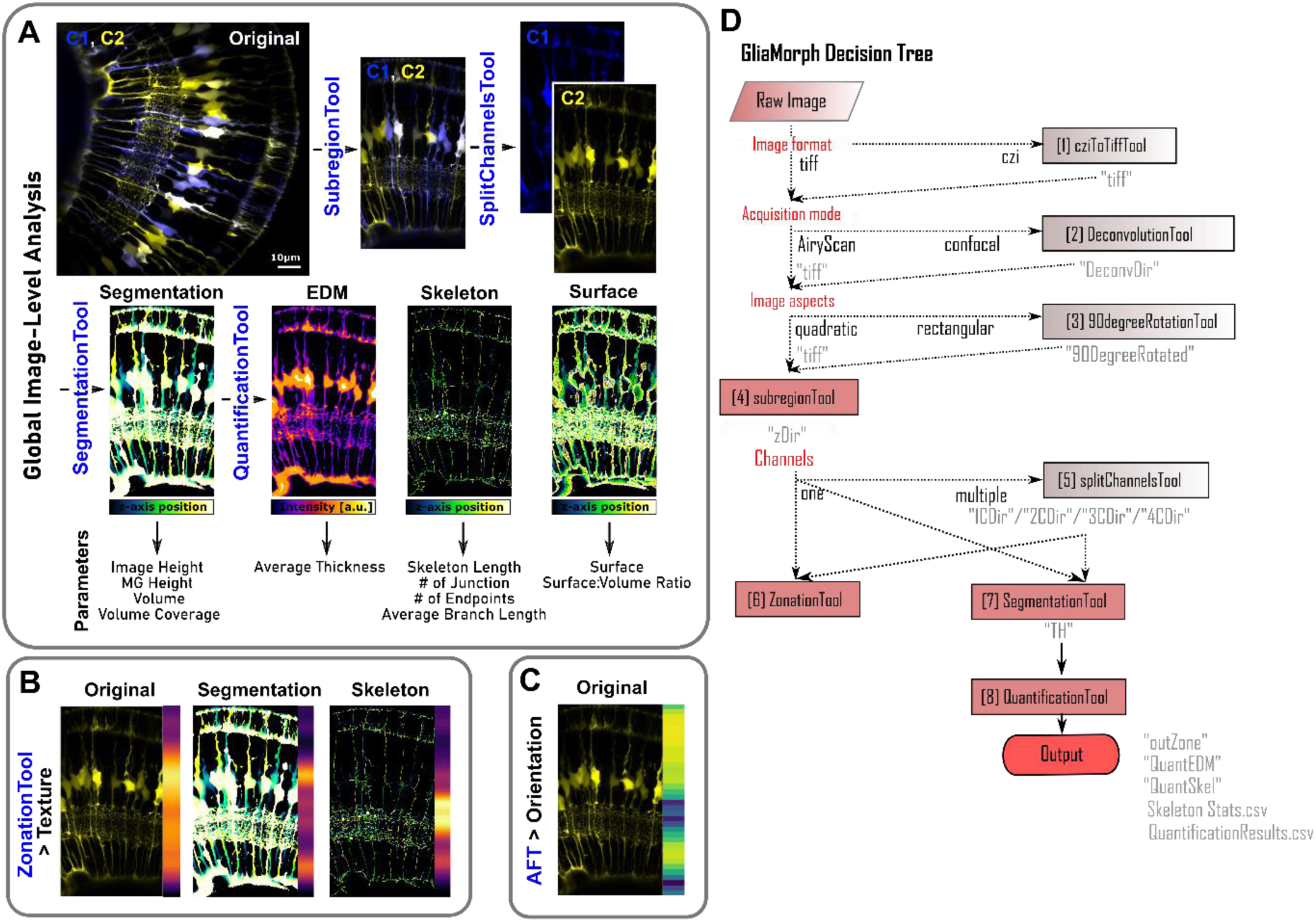
GliaMorph workflow overview. **(A)** Using global image-level measurements, 11 different parameters are quantified. **(B)** Using the zonationTool and applying it to original, segmented, and skeletonized data allows insights into apicobasal subcellular feature distributions. **(C)** Using Fourier Transformation based analysis allows the assessment of apicobasal subcellular orientation distributions. **(D)** As GliaMorph is modular in its application, workflow design is easy and flexible.

## Discussion

### The Importance of Imaging Data Quality and Standardization

We presented a comprehensive data analysis workflow to assess 3D MG morphology. We show that in-depth data understanding is crucial to analysing data in 3D. We performed benchmarking and troubleshooting using a variety of experimental approaches to identify key parameters that must be optimal/optimized for workflow validity. The better the data, the better the data analysis output. This is particularly important when assessing the apicobasal polarity found in MG, which translates to 5 subcellular domains that are biologically and computationally highly distinctive. To obtain easily comparable datasets, one should pay particular attention to the standardisation of position during image acquisition. For the samples we examined we acquired data in the ventro-temporal zone of the right eye (dorsally to the area temporalis, known as strike zone [38], or high acuity area). This standardization allowed comparability between samples. Also, image acquisition was standardized to be at the depth of the vascular inner optic circle with MG being parallel to the imaging plane. This enabled the complete tracing of MGs rather than analysis of intersections. However, standardization procedures might differ for samples at other ages, visualization techniques (e.g., different transgenics, antibodies, microscopes, etc.), or species.

To visualize data with a sub-diffraction resolution, we used AiryScan microscopy. We highlighted challenges in z-axis signal decay, and we suggested that the appropriate deconvolution is a requirement for fluorescence microscopy, which is in line with previous work [29], [39]. However, particularly for antibody staining, penetration depth is a limiting factor in image quality, and can only be addressed to a limited degree with computational processing. Again, these observations highlight that high input data quality is pivotal for quantitative image analysis.

As data are often acquired at different orientations, we achieved data comparability by employing semi-automatic rotation and 3D subregion selection. Standardization allowed for dimensionality reduction using the zonationTool to create 1D vectors from 3D data, enabling intuitive and quantitative insights into apical-basal polarity data. Lastly, employing standardized data, we used fibre-orientation assessment to analyse vertical-to-horizontal structures in our data.

Together, high-quality data, data understanding, and establishing image comparability allowed for data analysis and quantification on various levels.

### Biological Findings enabled by GliaMorph

GliaMorph allows for multi-dimensional glia analysis, enabling image-level as well as subcellular assessments that are robust, easy to use and adaptable.

For 3D morphological analysis, membrane markers generally showed better visualization vs. cytosolic markers. This was true in clonal and whole-transgenic analysis. However, when examining a membrane-marker injected into the reporter line *Tg(TP1bglob:VenusPest)^s940^*, occasional mislabelling of the former was observed. We believe this to be the case as VenusPest is a destabilized fluorophore with a rapid turnover, while GFP lingers longer. False labelling may label earlier born neurons, such as amacrine cells (**Fig. S6J)**. Additionally, even though glia shape serves as a readout of maturity [7] or cell damage [10], plasma membranes can suffer disruption, interfering with morphological analysis [10].

Our work further examines cell heterogeneity and compared whether individual cells represent collective measurements of cell populations. We show that single-cell and global image-level measurements are in agreement, but that batch effects might affect measurements. Using different analysis approaches, we performed global image-level, apicobasal texture, and apicobasal orientation measurements. Applying these different approaches to *cdh2* crispants, we show that *cdh2* loss leads to MG shape changes that were restricted to subregions. Applying apicobasal texture analysis to a mouse disease model, we show that there is an overall shape change as well as reactive cytoskeletal components in glaucoma.

### Robustness of Application

One aim for data analysis approaches should be robustness across users and data. Using acquisition standardization and automatic analysis ensures comparability between age-matched samples. This is exemplified when acquiring two datasets by two independent investigators and comparing MG volume and skeleton as readout. This showed neither a significant difference nor bias for both measured parameters. Thus, even if data are acquired in different samples and by different people, age-matched analysis is possible (**Fig. S9A, B**; volume p=0.9211; skeleton p=0.8460). Similarly, analysis of the same dataset, but by different experimenters does not bias/change the parameters analysed by GliaMorph (**Fig. S9C,D**; volume p=0.1934; skeleton p=0.7363). Additionally, we here presented data acquired by markedly different approaches, varying experimenters, transgenics/antibodies, wholemount/sections, different microscopes, and zebrafish/mice. Being able to perform multi-dimensional data analysis and detect subtle differences that might be otherwise overlooked using visual or manual assessments, we are confident that future work can use GliaMorph to aid decipher disease courses and answer questions such as “Does glial shape alteration precede neuronal dysfunction in neurodegenerative disease?”. Currently, GliaMorph is designed to perform on single-timepoint acquisitions, but this can easily be adapted if time-points are saved as individual stacks. Attention must be however paid if motion correction is required as this has to be applied prior to analysis. Thus, utilizing computational tools there is an exciting future ahead.

### Applicability to MG in other Species and other cell types

Here we show that GliaMorph can detect sub-cellular MG morphology changes in the retinas of crispant and disease models. As GliaMorph is modular it allows for adaptability to other MG visualization techniques. This could be other transgenic reporter lines or antibody staining, but also data from different species, and potentially other cell types, including other glial cells. The main bottleneck for transferability is the segmentation step, which extracts objects from the background. This segmentation can be performed in 2D (for a 3D stack this would mean slice-by-slice) or 3D (time-series data are traditionally segmented as 3D+t rather than 4D). Importantly, even if visualizing the seemingly same structure, if the visualization approach differs (e.g., different transgenic construct), segmentation is unlikely to be transferable between those [40]. This is because segmentation is influenced by a manifold of factors such as intensity, colour, texture, and in some cases connectivity. Additionally, data can be influenced by partial volume effects (*i.e.,* insufficient sampling frequencies), artifacts (motion artifacts, ring artifacts, autofluorescence) or noise due to sensors or electronics [41], [42]. Another consideration is the analysis of whole-mount intact tissues vs. cryo-sectioned tissue data, as the latter tends to be impacted by tissue rupture. To address some of these aspects, in recent years, machine learning has emerged as a meaningful approach for image segmentation, however, it often relies on having a ground-truth [43], [44], which is still widely lacking in biomedical image analysis.

Together, all steps – *except image segmentation* – of the GliaMorph toolkit are directly applicable to data other than the ones presented here. However, application to other data needs to come with the *caveat* that data often differ in sample preparation (e.g., different antigen retrievals, bleaching, or fixation) and image properties (e.g., autofluorescence). As mentioned, the main bottleneck is the segmentationTool (**Table 1**) which will require optimization for any data analysed other than the ones presented here. Most of the developed tools will be independent of input data properties such as microscope-type or cell features (e.g., microglia show a radial structure), see **table 1** for more detail. However, for all analyses, good image quality and standardization are key for meaningful quantifications.

**Table 1.**
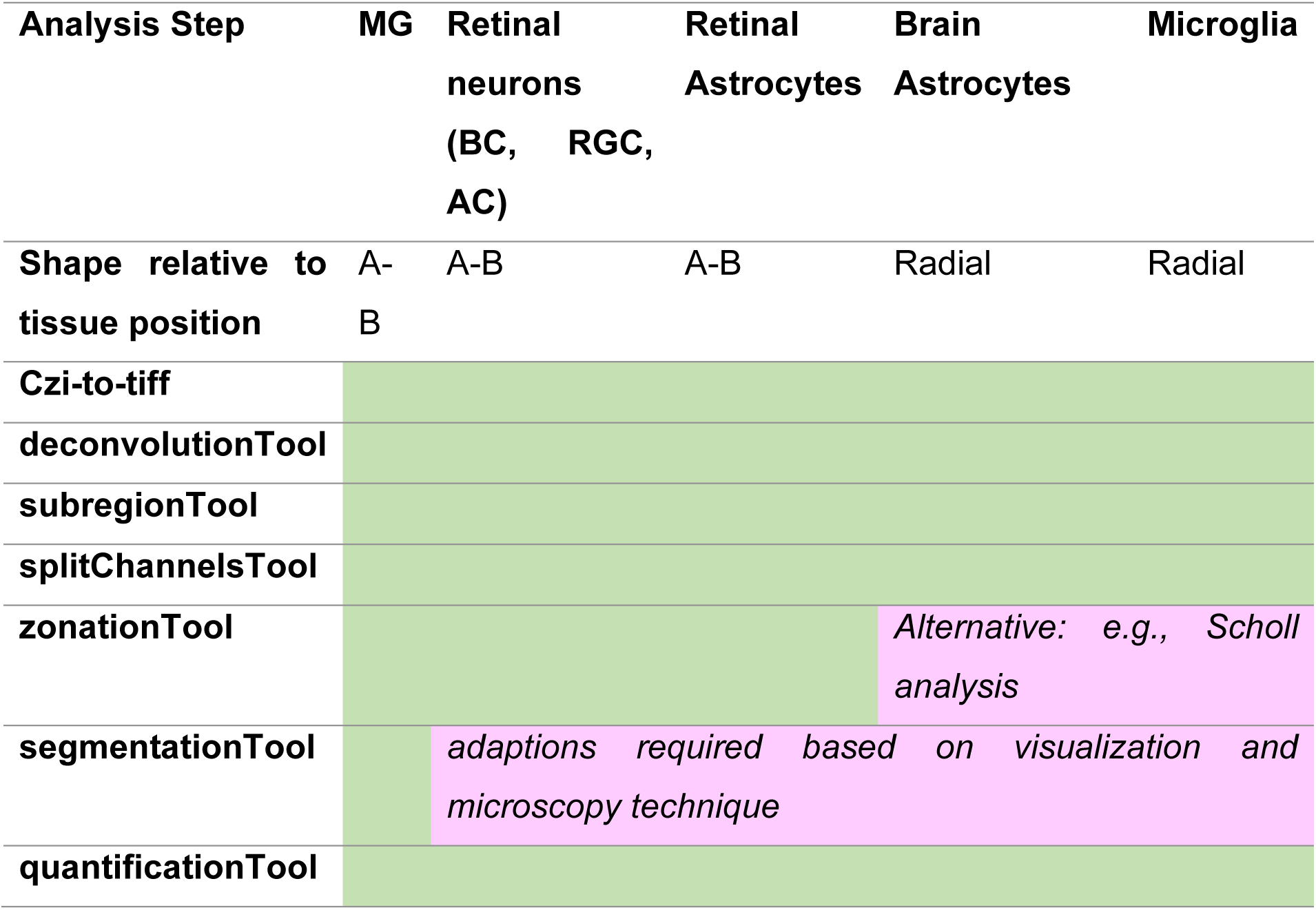
Generalizability of GliaMorph tools.

We here presented an in-depth analysis workflow to quantify 3D MG morphology. GliaMorph could be adapted to computationally automatically extract individual MG in segmented images to derive single-cell based measurements. This is challenging as cells are physically connected and while humans could potentially visually discern individual cells, computers often require *a priori* information or boundary conditions. On the other hand, unsupervized approaches, such as water-shedding [45], are limited in applicability to complex shape. However, to examine cell heterogeneity, computational identification of individual cells by cell separation is a crucial step to be examined in future work. For many studies, the molecular mechanisms regulating the establishment and maintenance of MG morphology remain elusive, due to the crude and/or challenging quantification in mutants. GliaMorph will provide a robust computational pipeline to analyse and quantify MG shape in genetic mutants or knockdowns of candidate genes [9], [37]. Importantly, as techniques develop, future work will also be looking at integrating multimodal data, such as shape analysis, calcium data for cell function, and overall animal behaviours. Ultimately, understanding biological mechanisms at the most fundamental level will be essential to understanding the processes observed in human vision pathology and how these could potentially be prevented with the aid of early predictive data analysis.

## Conclusion

We presented a comprehensive image understanding and analysis workflow, which is modular and open-source in application. Our presented work is an important benchmark study on how to understand and analyse retinal MG data.

## Material and Methods

### Zebrafish Handling and Husbandry

Experiments performed at UCL conformed to UK Home Office regulations and were performed under Home Office Project Licence PP2133797 held by RM. Maintenance of adult zebrafish in the fish facilities was conducted in Aquaneering tanks with a density of 5 animals per litre according to previously described husbandry standard protocols at 28°C with a 14:10 hours (h) light:dark cycle [46], [47]. Embryos, obtained from controlled pair- or group-mating, were incubated in E3 buffer (5mM NaCl, 0.17mM KCl, 0.33mM CaCl2, 0.33mM MgSO4) with/without methylene blue and 0.0045% 1-phenyl 2-thiourea (PTU) [48] applied between 6-24hpf and refreshed at a minimum of every 24h.

### Zebrafish Strains

To visualize Müller glia the following transgenics were used: *Tg(tp1bglob:VenusPest)^s940^* [30], *Tg(CSL:mCherry)^jh11^* (also known as *Tg(tp1bglob:hmgb1-mCherry)^jh11^* [49]). Retinal ganglion cells, photoreceptors, amacrine cells and horizontal cells were visualized with *Tg(ath5:gapRFP)^cu2^* (also known as *Tg(atoh7:gap43-mRFP1)^cu2^* [50]). Bipolar cells were visualized with *Tg(vsx1:GFP)^nns5^* [51]. Amacrine and horizontal cells were visualized with *Tg(ptf1a:dsRed)^ia6^* [52] and *Tg(ptf1a:cytGFP)* [53].

### Mouse Animal care

All animal husbandry was conducted according to the guidelines of the Canadian Council on Animal Care, using uOttawa ethical protocols OHRI-2856 and OHRI-3499. Animals were housed under specific pathogen-free conditions in standard isolation cages with enrichments. Animals were provided with food and water *ad libitum*. CD1 mice were obtained from Charles River Laboratories, and DBA2/J mice were obtained from Jackson Laboratories. Animals of both sexes, aged 335 days to 1 year, were used in this study.

### Constructs Generation

The expression constructs *pTol2-tp1bglob:eGFP-CAAX;cmlc2:eGFP* and *pTol2- tp1bglob:mCherry-CAAX;cmlc2:eGFP* were generated according to the Tol2Kit [54], using Multisite Gateway Technology. p5E-tp1 was a gift from Nathan Lawson (Addgene plasmid # 73585) [55] pME-mCherry-CAAX was a gift from Yi Feng’s group. pME-eGFP-CAAXp3E-polyA and pDestTol2CG2 vectors were obtained from the zebrafish Tol2Kit.

### Microinjections and Generation of Stable Lines

Microinjections were performed using a borosilicate glass capillary needle (World Precision Instruments, TW100F-4) connected to a Pneumatic Picopump injector (World Precision instruments). To generate mosaic labelling of MG, 6.5pg of *pTol2- tp1bglob:eGFP-CAAX;cmlc2:eGFP* or *pTol2-tp1bglob:mCherry-CAAX;cmlc2:GFP* was co-injected with 25pg of purified capped Tol2 transposase mRNA in a volume of 0.5nL, into one-cell-stage zebrafish embryos. To establish the *Tg(tp1bglob:eGFP-CAAX*) stable line, injected embryos were screened for GFP expression in the heart at 48-72 hpf. The F1 generation was screened for eGFP-CAAX expression to identify founder fish with germline integration and stable transmission to offspring. Positively identified F1 larvae were grown to adulthood and a stable transgenic line was established based on F2 generation with the strongest and most pervasive expression.

### Cdh2 CRISPR/Cas9 Injections

CRISPR/Cas9 technology was used to edit cadherin2 (*cdh2 or N-cadherin*), a gene known to affect MG morphology [37]. gRNA sequences were predicted using ChopChop design tool [56] (https://chopchop.cbu.uib.no). To disrupt cadherin2 function, gRNA1: 5’-AATGTTCCGTACGGTAGCGG-3’ and gRNA2: 5’-TAAACGATGTACCGTTCCGG-3’, both targeting exon 1, were picked to be simultaneously injected. Synthetic gRNAs were obtained from Merck in a two-part (crRNA:tracrRNA) format. gRNAs were assembled by mixing equimolar amounts of each crRNA and tracrRNA. gRNAs were then pooled and Cas9 protein (ThermoFisher Scientific, A36498) was added. For control injections, a solution without crRNA was prepared. Zebrafish eggs were injected into the yolk, before cell inflation, with 1nl solution containing 1200pg total gRNAs and 2000pg Cas9 protein.

### Fixation

Embryos were fixed with 4% paraformaldehyde (PFA; ThermoScientific, 28908) in phosphate buffer saline (PBS) for 2-4h at room temperature. Dehydration was performed by consecutive 5min washes with 25%, 50%, 75% in PBS and twice 100% methanol (MetOH). Samples were stored at -20°C in MetOH. Rehydration was performed in the reverse order and followed by 3x 5min washes in PBS. Fixation was conducted at 24, 48, 60, 72, 96, and 120hpf (staging based on anatomical features).

### Zebrafish Immunohistochemistry

Embryos were incubated in 1μg/ml of 4′,6-diamidino-2- 532 phenylindole (Roche, Cat# 10236276001) at RT for 10min and washed in PBS 1x. *Tg(TP1bglob:VenusPEST; - 5.5 ptf1a:DsRed)* embryos were fixed in 4% paraformaldehyde overnight at 4°C. Immunostaining was carried out on whole embryos as previously described [57], with heating in 150 mM Tris-HCl pH 9.0 at 70°C and subsequent acetone incubation. Embryos were then incubated with primary antibodies mouse anti-Gfap (zrf1; ZIRC - 1:100), rabbit anti-Rlbp1 (1:200, 15356-1-AP; Proteintech, Rosemont, Illinois, USA), mouse anti-Glutamine synthetase (1:200, mab302, Merck, Kenilworth, NJ, USA), followed with secondary antibodies Goat anti-mouse IgG Alexa Fluor® 647 (A21235; Thermo Fisher Scientific, Waltham, MA, USA) and Goat anti-rabbit IgG Alexa Fluor® 488 (A11008; Thermo Fisher Scientific, Waltham, MA, USA).

### Mouse Immunohistochemistry

Immunohistochemistry was performed essentially as described previously [58]. Briefly, eyes were harvested from mice euthanized by CO2 overexposure before cervical dislocation, and the corneas and lenses were removed. Eye cups were fixed with 4% paraformaldehyde/PBS for 10 minutes at room temperature. After cryoprotection overnight in 20% sucrose/PBS, retinas were cut into 18 µm sections. Primary antibodies were anti-Cralbp (Rlbp1; Fisher antibody MA1-813) and anti-Gfap (Millipore Sigma AB5804).

### Image Acquisition

Zebrafish images were acquired on the Zeiss LSM 900 with Airyscan 2 using a 40x water-immersion LD C-Apochromat (NA 1.1) or 63x oil-immersion Plan Apochromat (NA 1.4). Laser lines 405, 488, 561, 640 nm. Embedding was performed using glass-bottom dishes and 1% low-melting-point (LMP) agarose (Sigma, A9414); which was covered following solidification with E3.

Mouse data were acquired on a Zeiss LSM900 with Airyscan 2 using a Plan-Apochromat 63x/1.40 Oil DIC f/ELYRA.

***Sampling frequency*** was assessed for *Tg(TP1bglob:VenusPest)^s940^* using the Nyquist sampling rules (https://svi.nl/NyquistCalculator; *Eq. 1-3*).

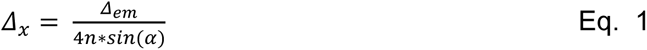

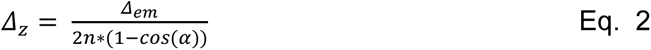

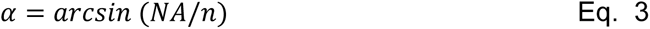

### Image Analysis

Images were analysed using open-source software Fiji [25]. Samples were excluded from analysis if image quality did not allow for reliable quantification (e.g., significant intra-plane movement, low CNR due to transgenic reporter weakness, unviable cell or animal health). Animals were allocated to treatment groups randomly without selection. Imaging and data analysis were performed unblinded to treatment allocation, often because the effect of treatment was easily deduced from the appearance of the micrograph. To overcome subjective bias, objective automized image analysis was applied where possible.

### Image understanding

Manual measurements of retina size and MG cell bodies were performed using a line region of interest (ROI; details see workflow document).

Manual measurements of intensity distributions were performed using a line ROI and plotting intensity values from maximum intensity projections (MIPs).

Contrast-to-Noise Ratio (CNR; Eq. 4) and Signal-to-Noise Ratio (SNR; Eq. 5) were quantified by the placement of 3-5μm circular ROI at slices of interest at the position of the cell body, protrusion, and endfoot for mean signal measurement, non-signal ROIs were placed inside the retina without cellular signal, and background ROI was placed outside the retina.

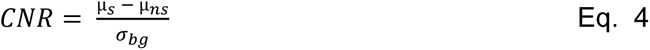

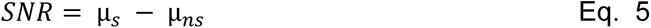

Where ***µs*** is the mean signal, ***µns*** is the mean non-signal, and ***σbg*** is the standard deviation of the background.

### Image Pre-processing

***“SubregionTool”***- Rotates images to align them along the y-axis, based on manual line ROI selection (start point at the MG endfeet and end point apically). Following the rotation, a bounding box is established with a 60 μm default width (x-axis) and box length is defined by the length of the manual line ROI with an extension of sigma (default 10 μm) to allow for the inclusion of the underlying vasculature and consideration of retinal curvature. The position of the bounding box is determined by the endpoint of the line ROI following rotation with the following potential cases: [1] x,y-defined box fits into the image; [2] box overfits on the right, meaning the box will start at point xT-bT, with xT being the image width and bT being the box width; [3] box overfits on the right, meaning the box will start at x=0; [4] box overfits on the bottom, meaning the output image height will be the length of the manual line ROI (error message “File X sigma could not be attached”, with X being the file name). Next, the image is cropped in x,y to this size saved, and the stack is reduced to the default 10 μm thickness. Should the stack not be thick enough an error message is produced: “File X not enough slices for substack”, with X being the file name.

***“90DegreeRotation”*** - Rotates images 90 degrees to the left or right before application of the *“SubregionTool”* and is required when images are rectangular rather than cubic, potentially resulting in artificial cropping.

N-tuple multi-labelling was established by image merging and false-colour LUTs application following the application of the *“SubregionTool”* to the respective embryos.

***“ZonationTool”*** – Following 8-bit conversion, MIPs (x, y-direction) were produced, images were transformed 90 degree from the left (Image > stack > reslice; **Fig. 3D**) and additional MIPs (z-direction) produced. For intensity plotting, image width can be scaled to 1920 voxels using bilinear interpolation to allow for comparability between different images. Line ROI was automatically placed starting from pixel 0 (left-most) to the image width, intensity profiles were measured and saved (one output file per input folder). One-voxel wise representation was assigned LUT Fire based on intensity and saved. Derived measurements were the total image height (*IN*) and retina height (*RN*), whereas retina height is based on sigma added with the SubregionTool.

**Table 1a.**
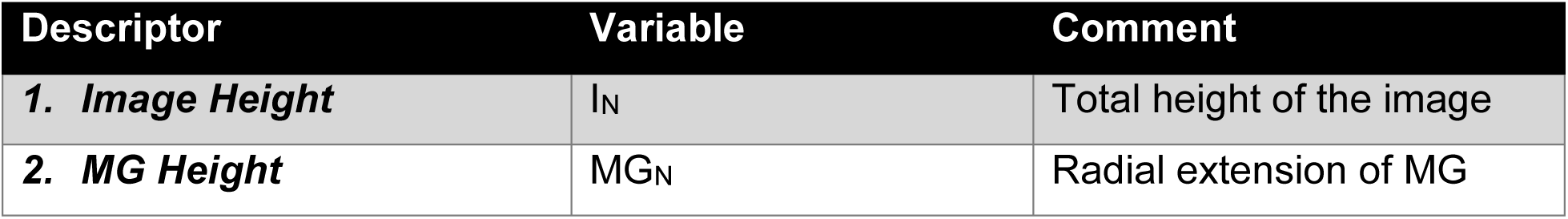
zonationTool measurements.

***AiryScan Processing***. AiryScan processing was conducted (independent of transgenic or staining) with Zeiss Zen Blue Software using 3D standard if not otherwise indicated in the text. Briefly, the processing is performed individually on images acquired with the 32 detectors, using a non-iterative linear Wiener deconvolution algorithm [59], [60].

As the processing is included in the Zeiss software system, there was no requirement to include additional steps into GliaMorph. However, we wanted to use this informed approach to establish the ideal processing before converting to GliaMorph. First, we examined the impact of AiryScan processing using the settings 2D standard, 3D standard, and 3D high (**Fig. S4A-D**). As expected, this showed 3D deconvolution to outperform 2D deconvolution, as assessed per reduced background (white arrowheads), and high deconvolution to result in increased structured noise and grains (black arrowheads). We next assessed this quantitatively using CNR measurements. This showed that 3D deconvolution indeed outperformed 2D processing and that reduction of 1μm z-stack step size to 0.19μm, increased the CNR drastically (**Fig. S4E,F**). As such, these parameters are feasible and ideal for MG morphological data in the zebrafish retina.

### “deconvolutionTool”

#### (1) Point-Spread-Function (PSF) Modelling

Theoretical PSF was modelled using analytical derivation based on Fraunhofer diffraction using the “Diffraction PSF 3D” Plugin (https://imagej.net/Deconvolution; https://www.optinav.info/Diffraction-PSF-3D.htm). The objective numerical aperture (NA) can be changed freely; the following standard suggestions were chosen: 20x Air NA 0.8, 40x Water NA 1.1, 63x Oil NA 1.4. Fluorophore wavelengths are entered manually for up to four channels.

#### (2) Confocal PSF Deconvolution

Deconvolution of the image ***y*** was performed with the Plugin DeconvolutionLab2 [61] (http://bigwww.epfl.ch/deconvolution/deconvolutionlab2/)

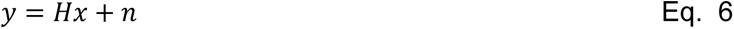

Where ***y*** is the data, ***H*** is the PSF matrix, ***x*** is the image, and ***n*** is the added noise component.

The following deconvolution algorithms were examined:

(a) *Regularized Inverse Filter* [62]:

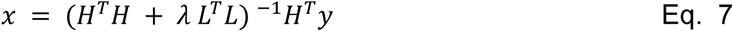

where ***L*** is the discretization of the Laplacian operator, ***^T^*** denotes the adjoint of L and H, and ***λ*** is the regularization factor (set to 1.000E-18).

(b) *Landweber* [63] *iterative deconvolution:*

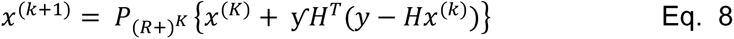

where ***P(R ^K^ = max(x,0)*** is the component-wise projection, the number of iterations ***Miter*** (set to 15), and step size parameter ***ƴ*** (set to 1.5).

(c) *Fast Iterative Shrinkage Thresholding* [64] *with the cost function:*

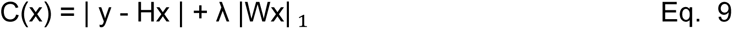

where ***W*** represents a wavelet transform and ***λ*** is the regularization factor (set to 1.000E-18).

(d) *Bounded-Variables Least Squares, also known as Spark-Parker algorithm* [65]*, minimizing a least-squares cost function (variables constraint):*

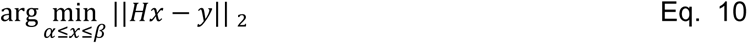

Where ***y*** is the vector of response variables, ***||.||2*** denotes the Euclidean norm, and ***α≤x≤β*** denote the upper and lower bounds, respectively.

(e) *Richardson-Lucy* [66], [67] *with the assumption of Poisson noise and the cost function:*

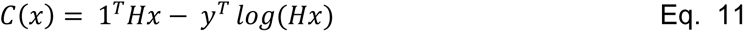

with ***Miter*** (set to 1 or 5).

To quantitatively assess image quality, contrast-to-noise ratio (CNR) was quantified. To account for subcellular differences (**Fig. S1A**) CNR was measured in several MG subdomains (MG cell bodies, protrusions, and endfeet). When examining the impact of RL iteration numbers visually, 1 iteration was found to preserve MG IPL protrusions (unfilled arrowhead **Fig. S3**) but not fully remove retina auto-fluorescence/background (green arrowhead), while 5 iterations did not fully preserve IPL protrusions but removed the observed background. This was confirmed when examining image intensities (**Fig. S3H**). When quantifying CNR levels, 5 iterations delivered higher CNR outcomes than 1 iteration (**Fig. S3I-K**). However, considering the loss of detail in IPL protrusions with 5 iterations, 1 iteration was used subsequently.

#### (3) User choice summary

The above was then integrated into the *deconvolutionTool* with user choices as follows: single-/multi-channel, selection of fluorophores by manual input of wavelength (nm) for up to 4 channels, NA input, and non-/existing PSF file.

### Segmentation

#### [A] Zebrafish Tg(TP1bglob:VenusPest)^s940^ and Tg(CSL:mCherry)^jh11^

Bleach correction was performed using “Simple Ratio Method” with background 0 [68]. Following, images were converted to 8-bit to allow for the following steps. Smoothing was performed using a 3D median filter with a radius of 2 voxels [69]. Segmentation was conducted after the selection of the plane at the middle of the stack with *Otsu thresholding* [70] *(histogram-derived):*

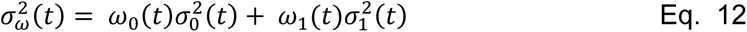

where ***ω_0_*** and ***ω_1_*** are the probabilities of the two classes to be separated with the threshold ***(t)***, calculated from the histogram, and ***σ_0_*** and ***σ_1_*** are the class variances, respectively. For post-processing, 3D median filtering with a radius of 2 voxels and binarization were conducted for surface smoothing. Additionally, MorpholibJ “Keep Largest Region” from the IJPB-plugins was used [71].

#### [B] Zebrafish Tg(TP1bglob: TP1bglob:eGFP-CAAX) ^u911^

Prior to segmentation, the “3D Edge and Symmetry Filter” was applied (settings: “alpha=0.500 compute_symmetry radius=10 normalization=10 scaling=2 improved”) [72]. As [A] above, with post-processing as follows: 3D hole filling [72] instead of surface smoothing.

### Quantification

All quantifications were performed using segmented 3D stacks as input.

To quantify the number of MG (**NN**), three approaches were tested (**a**) automatic ROI selection and MG counting, (**b**) semi-automatic ROI selection and automatic MG counting, and (**c**) manual measurements (taken as gold-standard). Briefly, to automatically select ROI, the ZonationTool was applied to the segmented 3D stacks to extract the MG cell body position per image, while for the semi-automatic approach, one rectangular ROI was drawn per experimental group (this assumes that animals are age-matched and comparable within the group). For automatic cell counting, stacks were cropped to ROI size in x,y. Then “Distance Transform Watershed 3D” [71] was applied to separate cell bodies, this was followed by “3D Simple Segmentation” [72] for binarization, and the BoneJ “Particle Analyser” [73] to quantify the number of cells.

MG volume (***VN***) was quantified as the number of object voxels (zero value) in histogram, multiplied by voxel size, while volume coverage (***VCN*** [%]) was measured as percentage of image voxels covered with MG object voxels. Surface (***SN***) were derived as surface/edge voxels following segmentation, and due to uncertainty in orientation (*i.e*. face, edge, or vertices) and normalization (*i.e.* voxel size differences between datasets) was given as volume a volume instead of area or number. Lastly, surface-to-volume (***S:VN***) ratio was derived as a ratio.

Centreline extraction was performed in segmented images using the Fiji “Skeletonize 2D/3D” Plugin, based on 3D thinning [74]. Centreline voxels (zero-valued in images) were quantified for total network length (***LN***) analysis by quantification of object voxels (zero value) in histogram.

The “Analyse Skeleton” Plugin in Fiji (Analyse > Skeleton > Analyse Skeleton 2D/3D [75]) was used to identify and measure the number of junctions (***JN***), End-Points (***EPN***), and average branch Length (***BLN***) (Analyse > Skeleton > Summarize Skeleton).

Euclidean Distance Maps (***EDMn***) of object voxel distance to the nearest background voxel were produced from binary segmented images using the Fiji plugin “Distance Map 3D”, which calculates distance in three-dimensional Euclidean space (*Eq.10*; Process > Binary > Distance Map in 3D [76]). To quantify thickness (***TN***), EDMs were multiplied with extracted skeletons, resulting in a 1D representation of vessel radii as represented by intensity of voxels (see [77]).

**Table 2:** Quantified parameters.

| Feature | Variable | Unit | Description |
| --- | --- | --- | --- |
| **Image Height** | $I_N$ | [$\mu\text{m}$] | Total height of the image |
| **MG Height** | $MG_N$ | [$\mu\text{m}$] | Radial extension of MG |
| **Number/Count** | $N_N$ | | Number of objects in ROI |
| **Volume** | $V_N$ | [$\mu\text{m}^3$] | Volume of object voxel, derived after segmentation |
| **Volume Coverage** | $VC_N$ | [%] | Percentage of image volume covered with MG (lowest = 0; highest = 100) |
| **Surface** | $S_N$ | [$\mu\text{m}^3$] | Number of object surface voxels, derived after segmentation (given in [$\mu\text{m}^3$] for comparability between experiments) |
| **Surface:Volume Ratio** | $S:V_N$ | | Ratio of surface to volume (lowest = 0; highest = 1) |
| **Thickness** | $T_N$ | | Distance from the local centreline to the corresponding surface |
| **Skeleton length** | $L_N$ | [$\mu\text{m}$] | Skeleton voxels (given in [$\mu\text{m}$] for comparability between experiments) |
| **# of Junction** | $J_N$ | | Number of points where 2/more sub-branches branch off |
| **# of Endpoints** | $EP_N$ | | Number of blind-ended object points |
| **Average Branch Length** | $BL$ | [$\mu\text{m}$] | Average length of individual skeleton branches |

### Fourier Transformation Analysis

To obtain information on MG alignment, AFT, a previously published open-source tool for evaluating the alignment of fibrillar structures across different length scales, was used [35]. This tool allows for local evaluation of alignment by exploiting the representation of small tiles of the image in the Fourier space. We achieved alignment analysis of MG by building on the AFT workflow as follows. We used (1) the SubregionTool to produce equally sized tiff MIPs, (2) “*CheckImageSize.ijm*” to identify the largest image, (3) “*NormalizeImageSizes.ijm*” to make data comparable by top-to-top alignment and attachment of black voxels to achieve image height similarity, using the largest image as standard – this step also includes an image enhancement step. We then applied (4) AFT only to non-black/non-background voxels (parameters: window size 100 px, overlap 50%, neighbour radius 3 vectors (*i.e., the neighbourhood of* 7 vectors); save output images – yes; apply filtering – yes; masking – 0, ignore blank spaces – 1, ignore isotropic regions – 0; mean background 5), and we produced (5) line-based averages along the apicobasal axis (i.e. use x averages, and iterate over y-axis).

#### Data representation

To visualize data, maximum intensity projections (MIP) were generated; intensity inversion was applied as appropriate to give the clearest rendering of relevant structures. Depth-coding was performed with the “Temporal-Color Coder” by Kota Miura (https://imagej.net/plugins/temporal-color-code).

### Statistical analysis

Normality of data was tested using D’Agostino-Pearson omnibus test. Statistical analysis of normally distributed data was performed using a One-way ANOVA to compare multiple groups or Student’s t-test to compare two groups. Non-normally distributed data were analysed with a Kruskal-Wallis test to compare multiple groups or Mann-Whitney test to compare two groups. Analysis was performed in GraphPad Prism Version 9 (GraphPad Software, La Jolla California USA). P values are indicated as follows: p<0.05 *, p<0.01 **, p<0.001 ***, p<0.0001 ****. All data were acquired as experimental repeats rather than replicates. Based on the mean/SD of the data in the examined groups in the assays used, post hoc power calculations have shown that these assays have at least 80% power to detect an effect size of 30% difference between groups when group sizes are 12/group (alpha = 0.05). Image representation was performed using Inkscape (https://www.inkscape.org). All data are available on request.

## Code and data availability

- **Code**: https://github.com/ElisabethKugler/GliaMorph
- **Example data for #GliaMorph Protocol:** 10.5281/zenodo.5747597
- **Minimum Example Data**: 10.5281/zenodo.5735442
- **Double-transgenic Müller Glia Data at 3dpf and 5dpf Zebrafish**: 10.5281/zenodo.5938758

## Acknowledgements.

We thank Robert Haase, Jonas Hartmann, Shanna Philip, and Eva-Maria Breitenbach for sharing ideas on how to improve the workflow and feedback on the manuscript. We thank the image.sc and Fiji community for ongoing support. The authors are grateful to Alessandro Felder from UCL Centre for Advanced Research Computing for guidance on RSE, code sharing, and code testing.

