## Supplementary Figures for "GliaMorph: A modular image analysis toolkit to quantify Müller glial cell morphology"

**Short title: Image-Driven Quantification of Retinal Glia Shape**

**Elisabeth Kugler[1§], Isabel Bravo[1], Xhuljana Durmishi[1], Stefania Marcotti[2], Sara Beqiri[1], Alicia Carrington[1], Brian M. Stramer[2], Pierre Mattar[3,4], and Ryan B. MacDonald[1§]**

1 Institute of Ophthalmology, University College London, 11-43 Bath St, Greater London EC1V 9EL.

2 Randall Centre for Cell & Molecular Biophysics, King's College London, New Hunt's House, London SE1 1UL.

3 Department of Cellular and Molecular Medicine, University of Ottawa, Ottawa, ON K1H 8M5, Canada.

4 Ottawa Hospital Research Institute (OHRI), Ottawa, ON K1H 8L6, Canada.

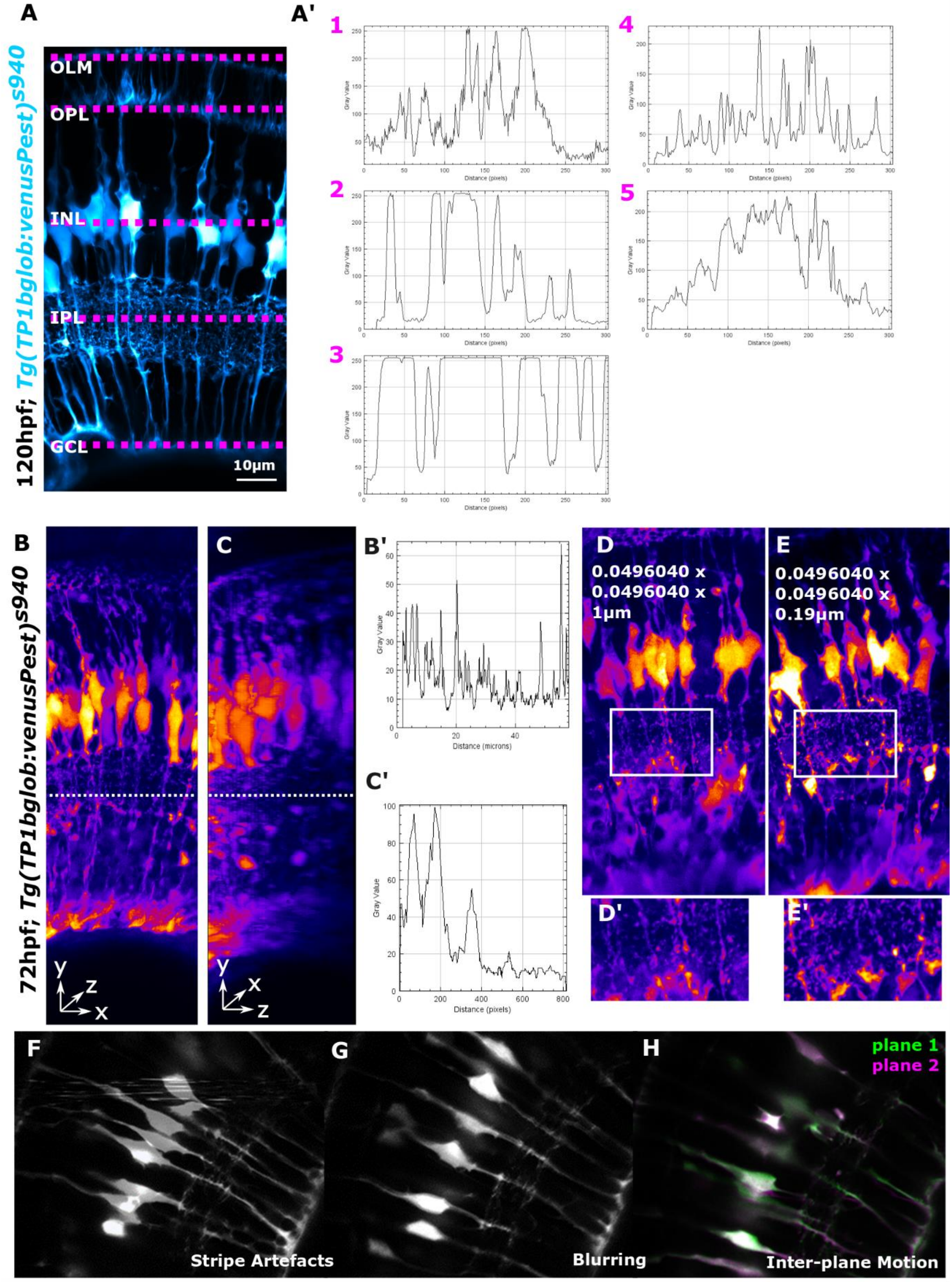


**Figure S1. Data Challenges.**

**(A)** MG subregions/zones (1-to-5) display different signal levels and patterns, as shown by intensity profiles in **A'**. **(B)** MIP of an image in the acquired direction with intensity profile **B'** along a region of interest (dotted line), and transformed to look at it laterally **(C)** showing signal decay in the z-axis **C'**. **(D)** MIP of sub-optimal (1µm) vs optimal (0.19µm) z-steps **E**. **(F-H)** Artefacts, such as stripe artefacts, blurring, and inter-plane motion, were observed (representative images).

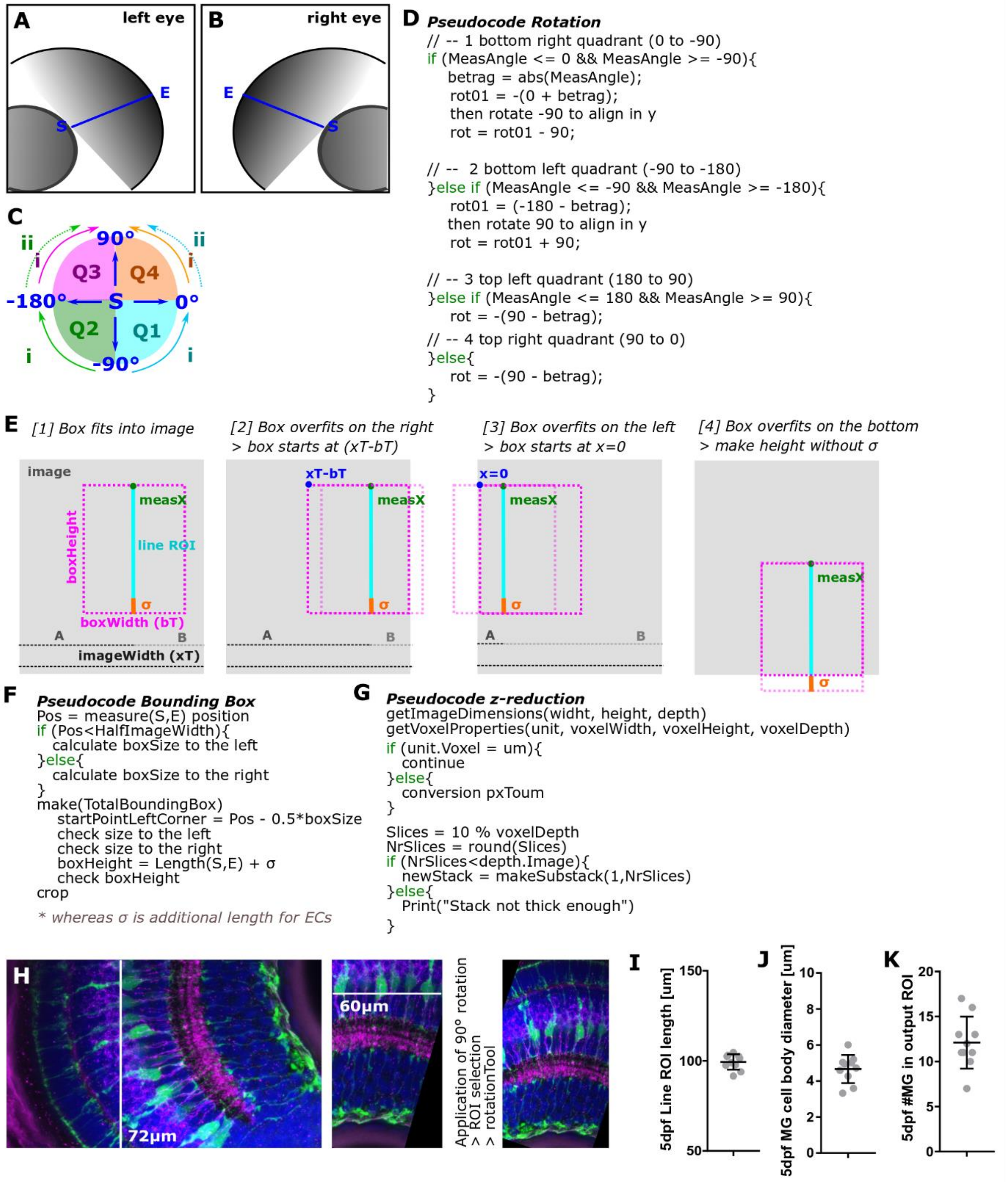


**Figure S2. Semi-automatic ROI selection.**

**(A-B)** The workflow applies to the left and right eyes, which is computationally an inverse problem (S – start; E - end). **(C)** Rotation is performed in a multi-step fashion, depending on the original image position. **(D)** Pseudocode for image rotation. **(E)** Bounding box starting points are determined based on the line ROI position. **(F)** Pseudocode for bounding box (x,y) establishment. **(G)** Pseudocode for z-reduction to establish stacks of the same depth. **(H)** For rectangular images, prior 90-degree rotation is suggested to avoid rotation-induced cropping. **(I)** The input line ROI determines the output image height, with the measurement of ROI length being an indicator for image similarity, such as an average 99.39 µm MG height at 120 hpf with a coefficient of variation (CoV) of 4.3% (n=10; N=3 experimental repeats; mean ± s.d.). To account for retinal curvature an additional section, called sigma (default 10 µm), is appended. **(J)** Z-depth is reduced to a default 10 µm, as cell bodies are an average of 4.6 µm in diameter at 120 hpf (n=10; N=3 experimental repeats; mean ± s.d.). **(K)** The output image width is set to a default 60 µm to include on average 12 MG at 120 hpf (n=10; N=3 experimental repeats; mean ± s.d.).

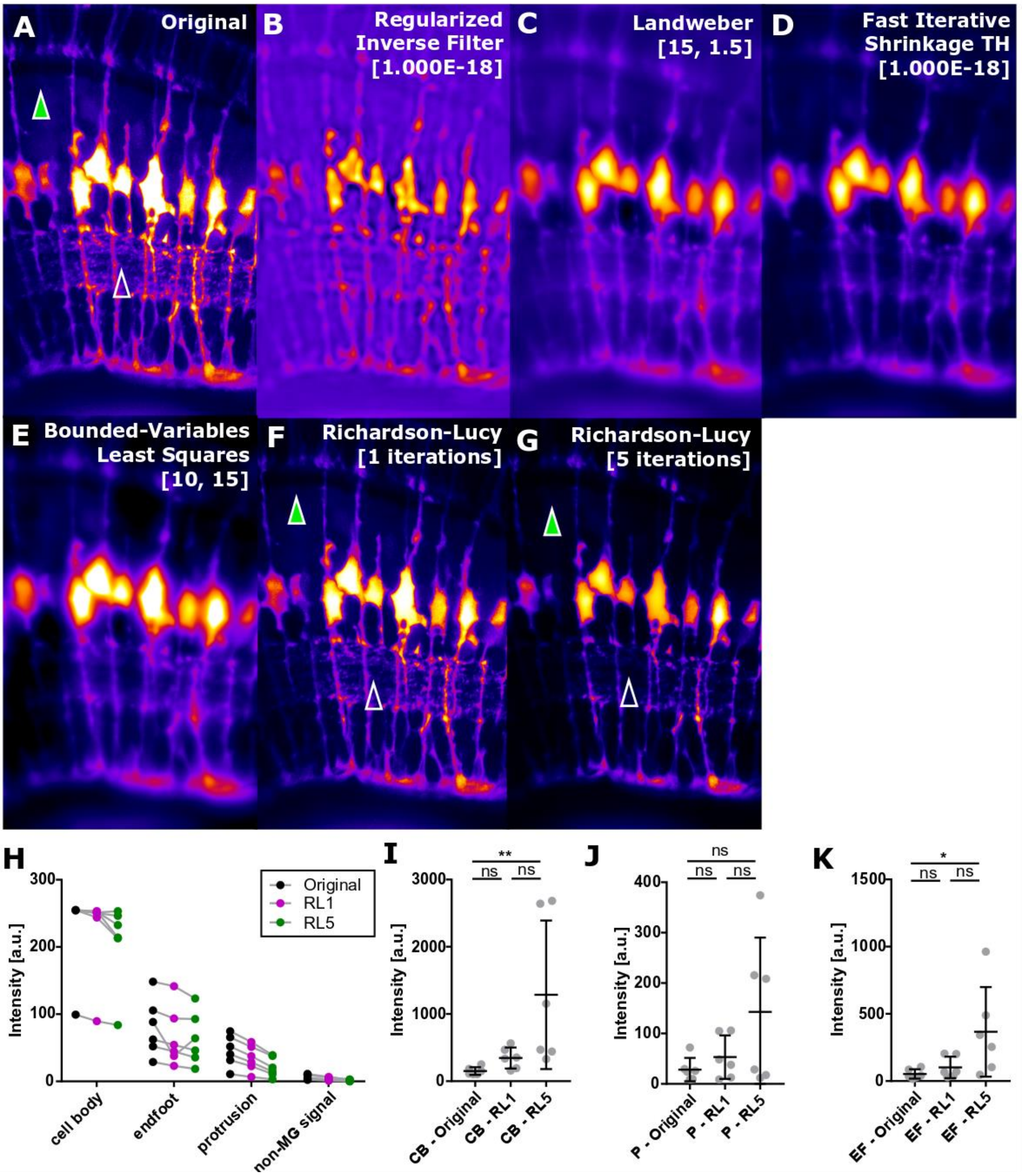


**Figure S3. Confocal data PSF Deconvolution.**

**(A)** Original image MIP (LUT Fire; green arrowhead – indicates non-MG background; unfilled arrowhead – MG IPL protrusions that could be lost with incorrect deconvolution). **(B)** Image after application of regularized inverse filter deconvolution. **(C)** Image after application of Landweber deconvolution. **(D)** Image after application of fast iterative shrinkage thresholding deconvolution. **(E)** Image after application of bounded least variables least-squares deconvolution. **F** Image after application of Richardson Lucy (RL) deconvolution with 1 iteration. **(G)** Image after application of RL deconvolution with 5 iterations. **(H)** Image intensity measurements in MG cell bodies, endfeet, protrusions, and non-MG background signal (n=6 120 hpf; black – original, magenta – following deconvolution with RL 1 iteration, green - following deconvolution with RL 5 iterations). **(I)** CNR quantification in cell bodies (CB; n=6 120 hpf; Kruskal-Wallis test; mean ± s.d.). **(J)** CNR quantification in protrusions (P; n=6 120 hpf; Kruskal-Wallis test; mean ± s.d.). **(K)** CNR quantification in endfeet (EF; n=6 120 hpf; Kruskal-Wallis test; mean ± s.d.).

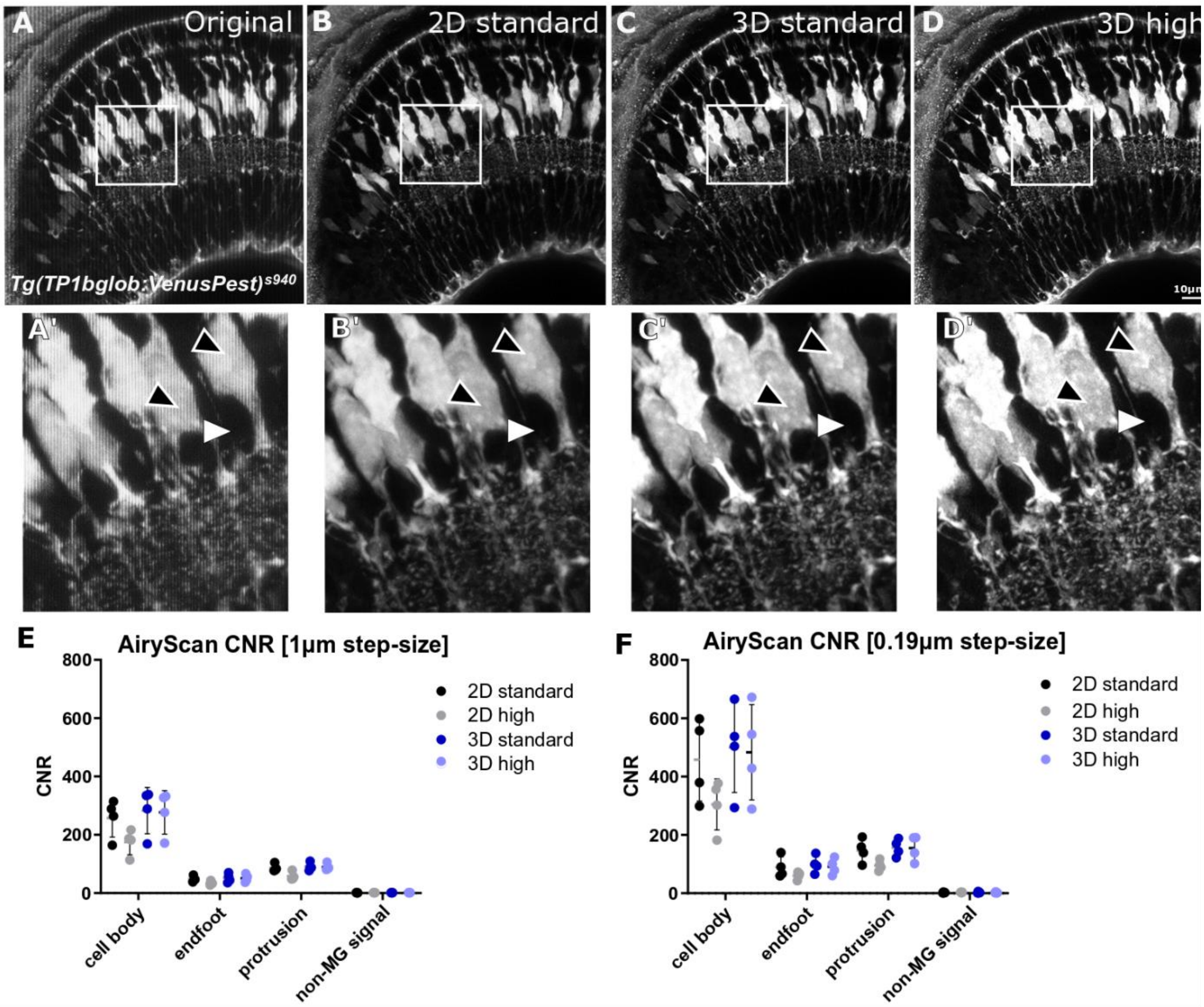


**Figure S4. Image AiryScan Processing.**

**(A)** MIP of the original image without pre-processing. **(B)** MIP of an image after 2D standard AiryScan deconvolution. **(C)** MIP of an image after 3D standard AiryScan deconvolution, showing decreased background signal (white arrowhead). **(D)** MIP of an image after 3D high AiryScan deconvolution, showing increased structures noise/grains (black arrowheads). **(A'-D')** Insets of A-D, respectively**. (E-F)** CNR measurements in MG cell bodies, endfeet, protrusions, and non-MG signal, processed in 2D/3D, standard/high, acquired with 1µm (E) and 0.19µm (F) z-stack step sizes (n=4 120 hpf; mean ± s.d.).

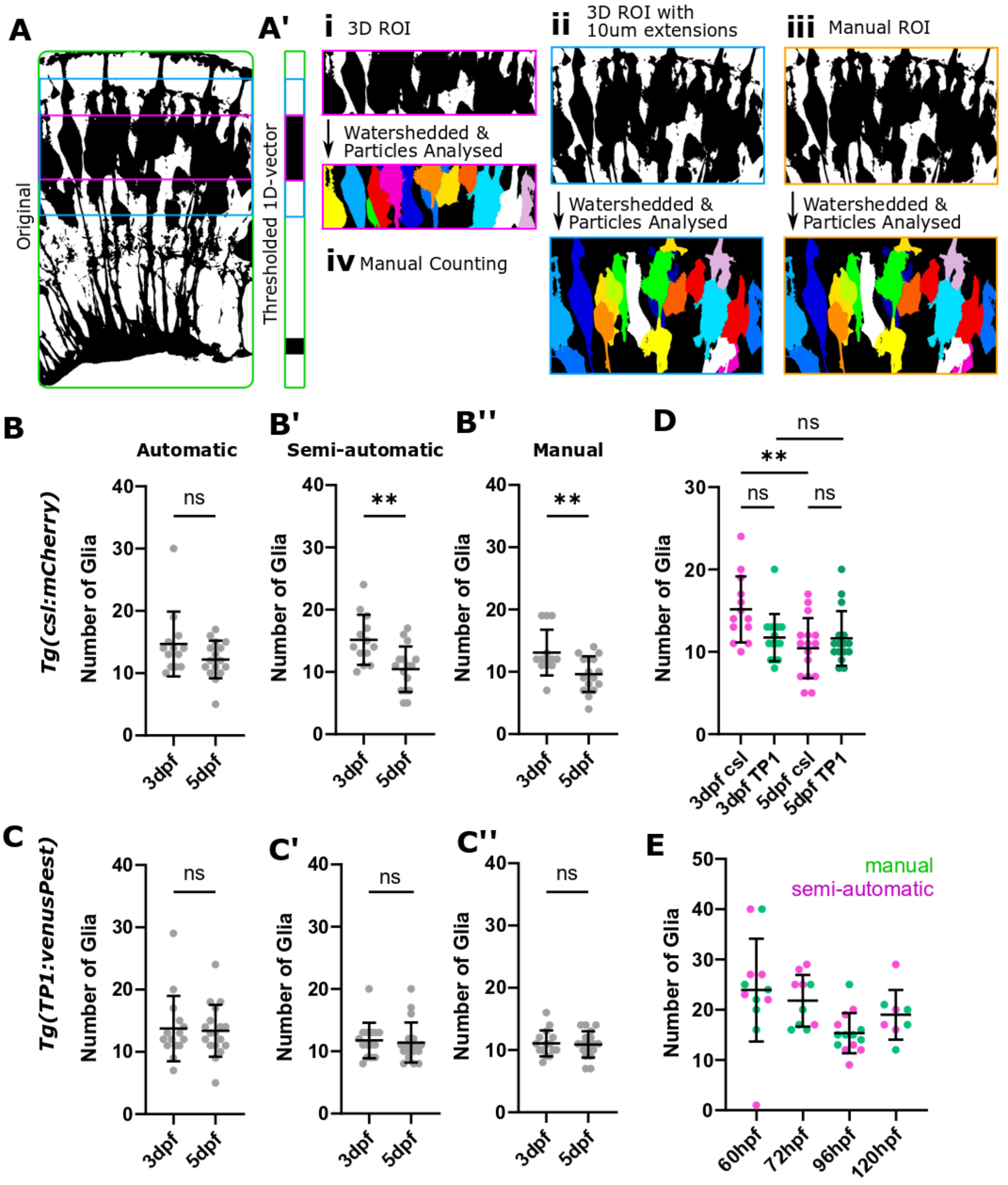


**Figure S5. Cell number counts are influenced by the visualization technique used.**

**(A)** Segmented 3D stacks were used to quantify the number of cells. **(A')** To automatically extract the retinal layer containing MG cell bodies, the ZonationTool was applied and then segmented (i). (ii) To allow for errors, the ROI from (i) was extended by 10µm in both directions. (iii) Alternatively, a semi-automatic approach was used, using one manually drawn ROI for each group. (iv) To compare measurements from the above, manual counting was used as gold-standard. **(B)** Comparison of measurement outcomes in *Tg(csl:mCherry)* (B p=0.0027, B' p=0.0027, B'' p = 0.0083; 3dpf n=13, 5dpf n=16; N=2; unpaired two-tailed Students' t-test; mean ± s.d.). **(C)** Comparison of measurement outcomes in *Tg(TP1:venusPest)*. This showed that semi-automatic and manual measurements were more sensitive than fully automated analysis. Also, this showed that the cell number decreased from 72-to-120 hpf in *Tg(csl:mCherry)* but not *Tg(TP1:venusPest)* (C p=0.8347, C' p=0.7500, C'' p=0.8103; 72 hpf n=13, 120 hpf n=16; N=2; unpaired two-tailed Students' t-test; mean ± s.d.). **(D)** Comparison of the glia number of both transgenics using semi-automatic measurements (mean ± s.d.). **(E)** Comparison of cell number measurements from 60-to-120hpf using semi-automatic and manual measurements (mean ± s.d.).

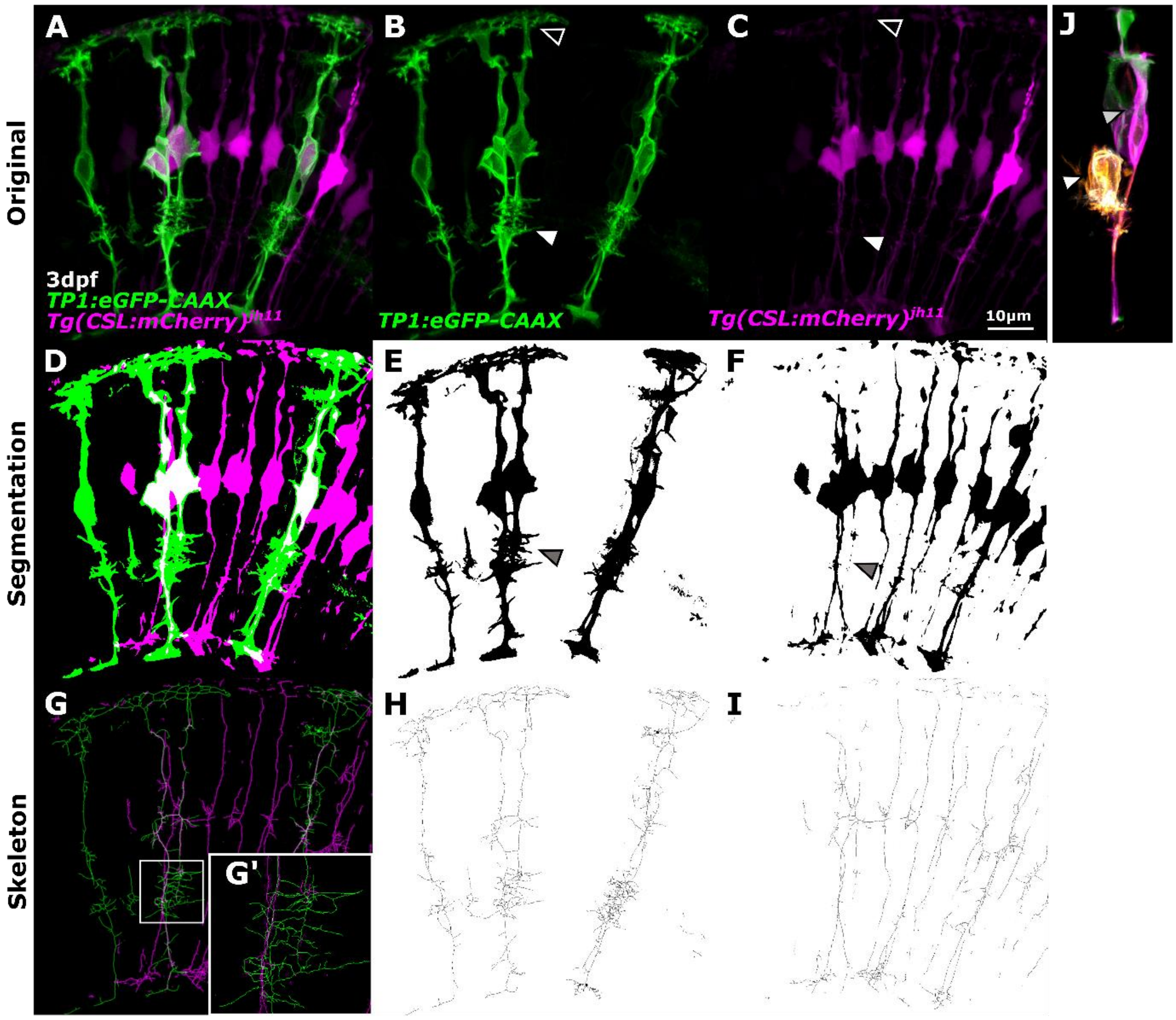


**Figure S6. MG membrane and cytosol segmentation show membrane signals to deliver better segmentation results.**

**(A-C)** MG cytosolic transgenic (magenta) with a mosaic expression of MG membrane marker (green). **(D-F)** Segmentation of membrane and cytosol markers. **(G-I)** Skeletonization of the segmented images. **(J)** Cell labelling can result in an unspecific signal, as seen by an individual MG (grey arrowhead) and amacrine cell (white arrowhead) being labelled.

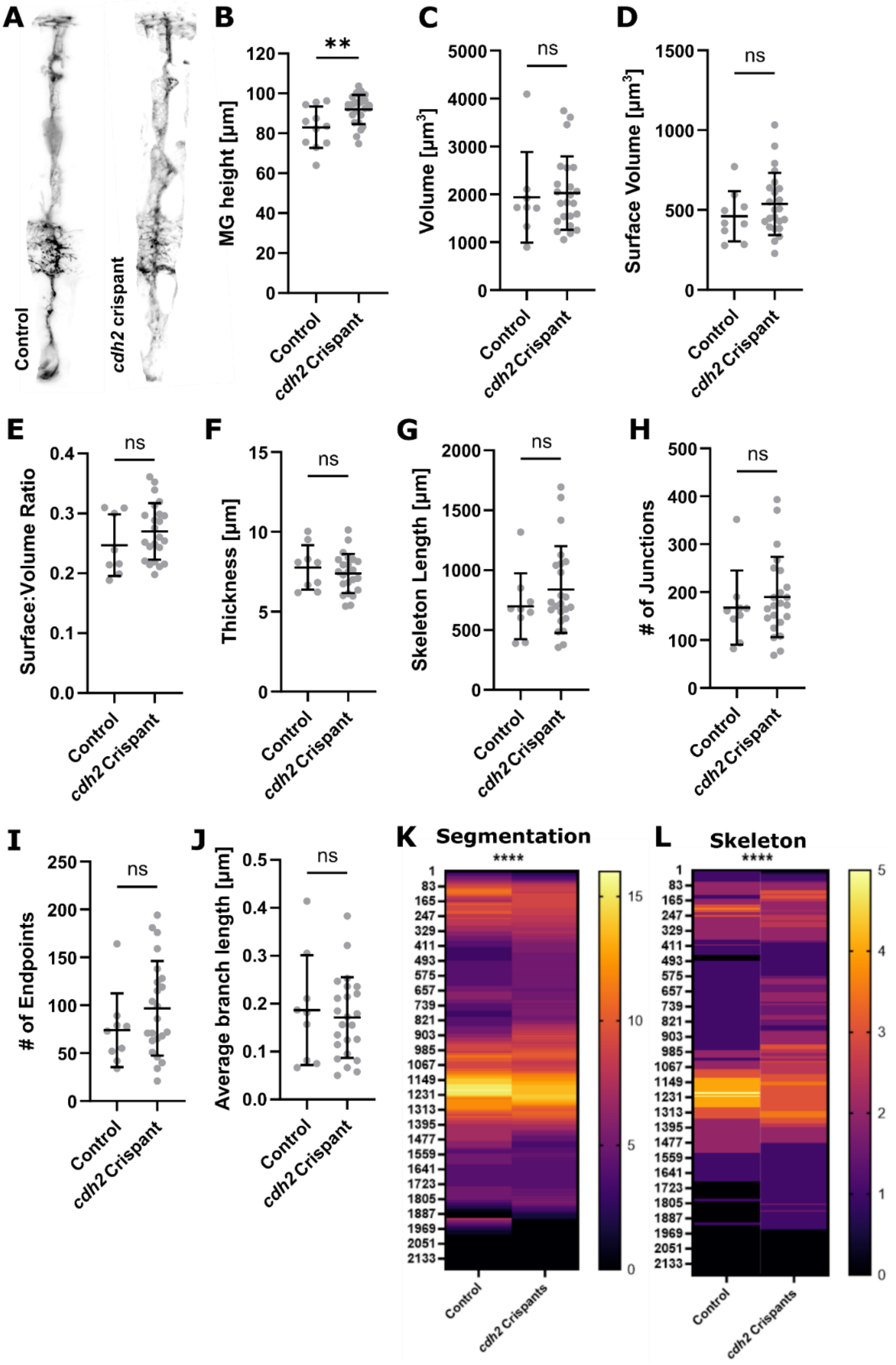

**Figure S7. Single-cell analysis of *cdh2* crispants.**

**(A)** Micrograph of control and *cdh2* crispants clones (representative images extracted from 3D stacks). **(B)** MG height was significantly increased in *cdh2* crispants (p=0.0049; control n=11 cells from 6 embryos, *cdh2* crispants n=26 cells from 12 embryos; N=2 experimental repeats; unpaired Students t-test; mean ± s.d.). **(C)** Volume was not significantly altered (p=0.7980; control n=11 cells from 6 embryos, *cdh2* crispants n=26 cells from 12 embryos; N=2 experimental repeats; unpaired Students t-test; mean ± s.d.). **(D)** Surface volume was not significantly altered (p=0.2984; control n=11 cells from 6 embryos, *cdh2* crispants n=26 cells from 12 embryos; N=2 experimental repeats; unpaired Students t-test; mean ± s.d.). **(E)** Surface-to-volume ratio was not significantly altered (p=0.2541; control n=11 cells from 6 embryos, *cdh2* crispants n=26 cells from 12 embryos; N=2 experimental repeats; unpaired Students t-test; mean ± s.d.). **(F)** Thickness was not significantly altered (p=0.4602; control n=11 cells from 6 embryos, *cdh2* crispants n=26 cells from 12 embryos; N=2 experimental repeats; unpaired Students t-test; mean ± s.d.). **(G)** Skeleton length was not significantly altered (p=0.3081; control n=11 cells from 6 embryos, *cdh2* crispants n=26 cells from 12 embryos; N=2 experimental repeats; unpaired Students t-test; mean ± s.d.). **(H)** Number of junctions was not significantly altered (p=0.4948; control n=11 cells from 6 embryos, *cdh2* crispants n=26 cells from 12 embryos; N=2 experimental repeats; unpaired Students t-test; mean ± s.d.). **(I)** Number of endpoints was not significantly altered (p=0.2252; control n=11 cells from 6 embryos, *cdh2* crispants n=26 cells from 12 embryos; N=2 experimental repeats; unpaired Students t-test; mean ± s.d.). **(J)** Average branch length was not significantly altered (p=0.2252; control n=11 cells from 6 embryos, *cdh2* crispants n=26 cells from 12 embryos; N=2 experimental repeats; unpaired Students t-test; mean ± s.d.). **(K)** Apicobasal texture plotting of segmented cell showed a statistically significant change (p<0.0001; control n=11 cells from 6 embryos, *cdh2* crispants n=26 cells from 12 embryos; N=2 experimental repeats; unpaired Students t-test; mean ± s.d.). **(L)** Apicobasal texture plotting of skeletonized cells showed a statistically significant change (p<0.0001; control n=11 cells from 6 embryos, *cdh2* crispants n=26 cells from 12 embryos; N=2 experimental repeats; unpaired Students t-test; mean ± s.d.).

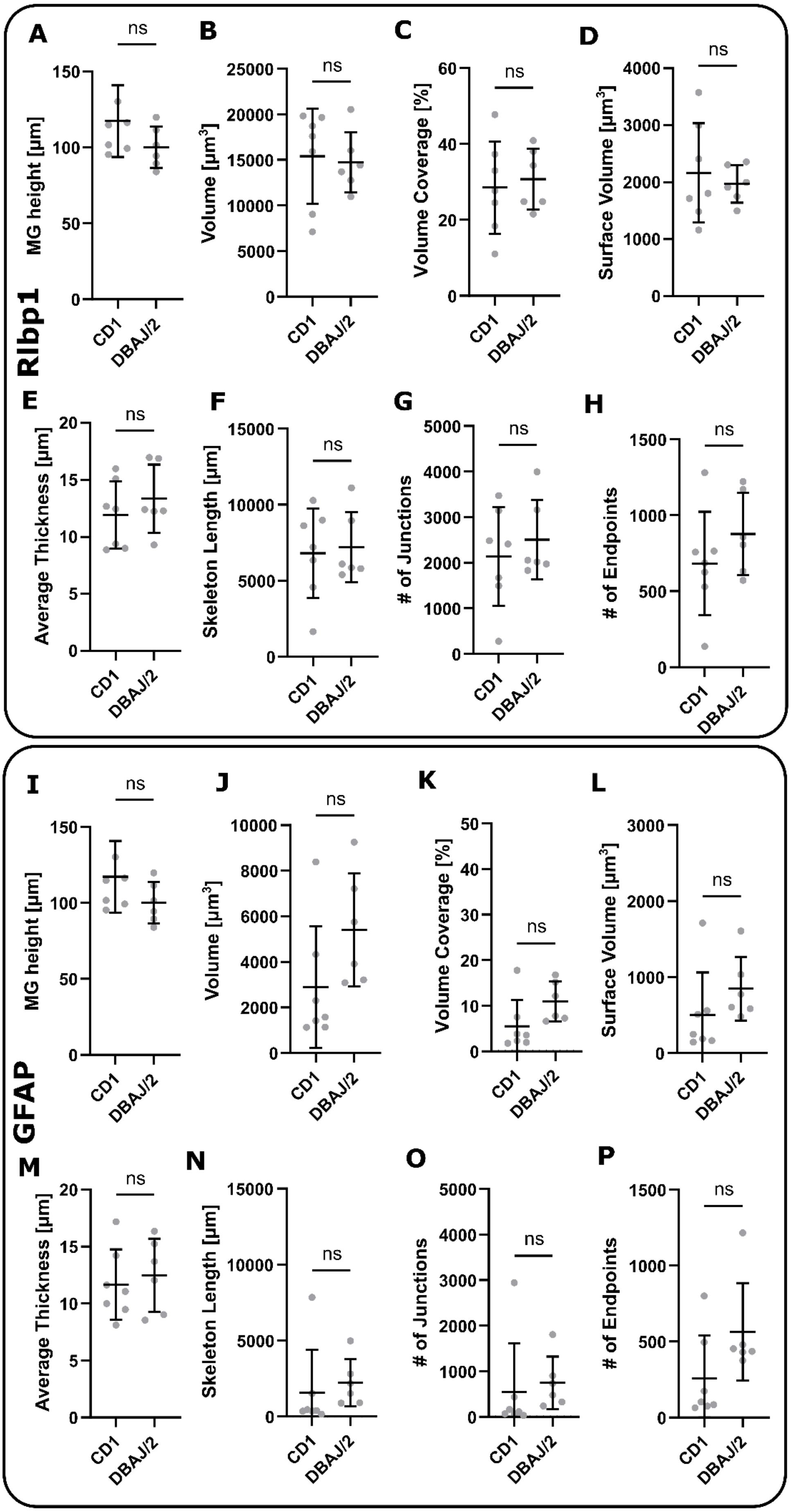
Rlbp1
GFAP
A
B
C
D
E
F
G
H
I
J
K
L
M
N
O
P
ns
MG height [µm]
Volume [µm³]
Volume Coverage [%]
Surface Volume [µm³]
Average Thickness [µm]
Skeleton Length [µm]
### of Junctions
### of Endpoints
CD1
DBAJ/2

138

**Figure S8. Mouse Glaucoma Model Quantification using the GliaMorph toolkit.** **(A-H)** Quantification of Rlbp1 data showed no statistically significant difference between CD1 controls and DBA/J2 (n=7 stacks from 3 mice each; MG height p=0.1375; Volume p=0.7308; Volume coverage p=0.6282; Surface volume p>0.9999; Average thickness p=0.6282; Skeleton length p=0.9452; Number of junctions p=0.6282; Number of endpoints p=0.2949; Mann-Whitney test). **(I-P)** Quantification of GFAP data showed no statistically significant difference between CD1 controls and DBA/J2 (n=7 stacks from 3 mice each; MG height p=0.1375; Volume p=0.0734; Volume coverage p=0.0734; Surface volume p=0.0734; Average thickness p=0.7308; Skeleton length p=0.0734; Number of junctions p=0.0734; Number of endpoints p=0.1375; Mann-Whitney test).

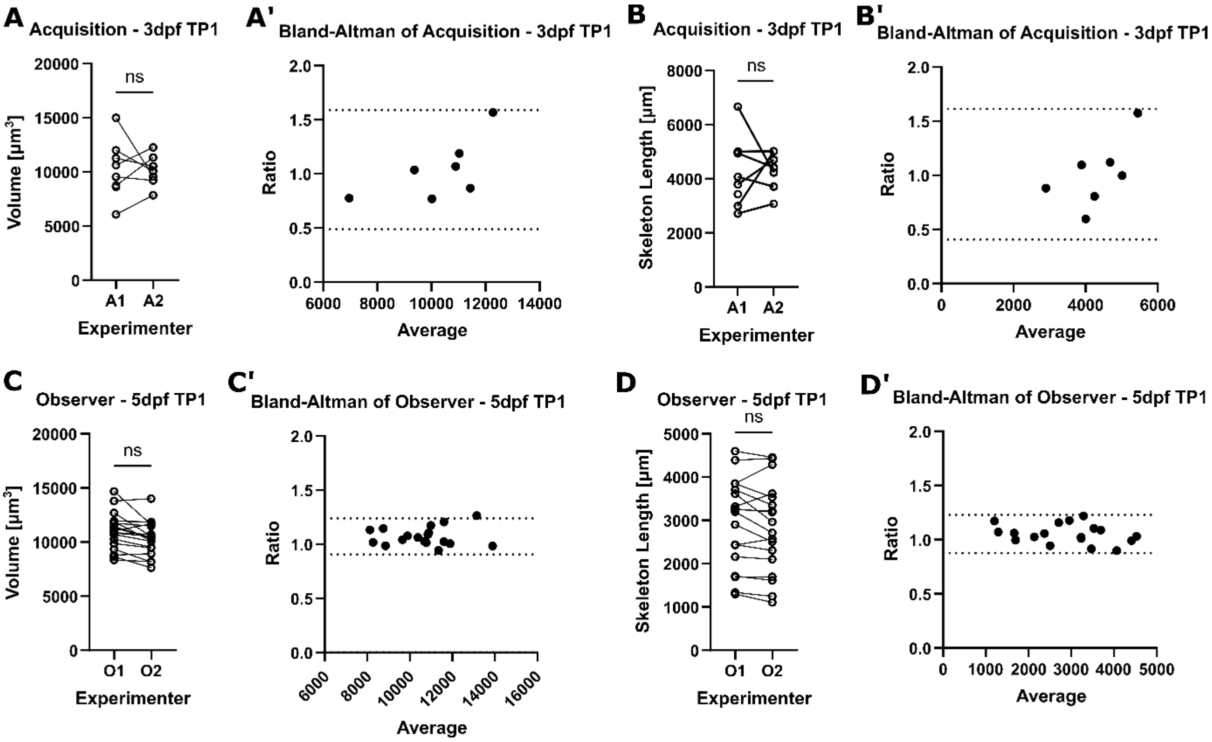


**Figure S9. Robustness of Analysis Workflow.**

**(A, A')** Comparing the analysis outcomes using MG volume from data acquired by two different people showed no statistically significant difference (p=0.9211 unpaired t-test) and no bias measured by the Bland-Altman ratio (1.039; n=8; N=1). **(B, B')** Comparing the analysis outcomes using skeleton length from data acquired by two different people showed no statistically significant difference (p= 0.8460) unpaired t-test) and no bias was measured by the Bland-Altman ratio (1.010; n=8; N=1). **(C, C')** Comparing the analysis outcomes using MG volume analysed independently by two people showed no significant difference (p=0.1934 unpaired t-test) and no bias measured by Bland-Altman ratio (1.073; n=18; N=2). **(D, D')** Comparing the analysis outcomes using MG skeleton length analysed independently by two people showed no significant difference (p= 0.7363 unpaired t-test) and no bias measured by Bland-Altman ratio (1.053; n=18; N=2).
